# From Diverse Prior Knowledge to Mechanistic Causal Network Using PSoup: A Case Study in Shoot Branching

**DOI:** 10.64898/2026.08.07.743620

**Authors:** Christos Mitsanis, Nicole Fortuna, Christine Beveridge

## Abstract

Mechanistic models of plant regulatory networks typically require extensive parameterization, limiting their generalisation and scalability. Here we present a parameter-free, topology-driven model of shoot branching that predicts phenotypic outcomes from network structure alone. We constructed a signed, directed causal network by distilling regulatory relationships from the published literature spanning many laboratories, species, years, data types, and methodological frameworks. This extracted the essential logic of the system, consistent with developmental-biological reasoning and anchored in empirical evidence. Using PSoup, which automatically translates network topology into algebraic equations, the model propagates information across the network and predicts the qualitative direction of change relative to a defined baseline, mirroring the comparative framework of biological experiments. The pipeline, from network construction through automated equation generation to prediction, is transparent and reproducible. Trained against branching phenotype data with 78 diverse perturbations spanning genetic mutations and hormone treatments, the model achieved 86% accuracy in predicting branching direction. On an independent test set of 84 perturbations measuring bud release and gene expression at nodes not used during training, accuracy reached 75%. The approach highlighted deficiencies in our understanding of the topology of the network around SMXL 6/7/8 and ABA nodes. Other errors came mainly from modelling choices, such as the threshold for scoring a node as changed relative to baseline. Beyond shoot branching, this work demonstrates a general strategy for synthesizing biological knowledge into validated predictive networks, providing a foundation for both applied breeding and the advancement of fundamental biology.

## Introduction

Shoot branching is regulated by a complex network that has long been studied at both fundamental (Beveridge et al., 2023) and agronomic levels, with impacts in agriculture (Alam et al., 2014; Borrell et al., 2022; Hammer et al., 2020) and forestry (Kint *et al*., 2010). Given its broad importance, a comprehensive model of shoot branching would be highly beneficial for agronomists, breeders, geneticists, plant physiologists and interdisciplinary collaborations within these fields (Hammer et al., 2006). Nevertheless, a comprehensive model encompassing the molecular physiology and genetics underpinning shoot branching has yet to be developed. Although incorporation of molecular networks into predictive models could theoretically contribute to better breeding and therefore contribute directly to crop improvement (Powell *et al*., 2026), such incorporation is challenged by lack of technologies to convert prior knowledge of complex networks, like shoot branching, into an appropriate model.

Such a model would, at a minimum, need to integrate information on the roles and interactions of the whole regulatory network, including genes and hormones such as auxin, cytokinin, strigolactones, abscisic acid, and gibberellins, as well as sugar and light quality (Beveridge *et al.,* 2023; Doidy *et al.,* 2024; Evers *et al.,* 2006; Leduc *et al.,* 2014). To date, existing models either partially cover some of these components (Bertheloot et al., 2020) or focus specifically on auxin flux according to canalization theory (Bennett *et al*., 2014; Prusinkiewicz *et al*., 2009; Runions *et al*., 2014) or light quality (Evers et al., 2006). Each of these models focused only on specific sections of the branching regulatory network and did not integrate processes occurring at multiple levels of organization in the hierarchy of subcellular to whole plant and environment. This focus of modelling only a specific part of a network is often accounted in modelling of biological systems.

Extensive knowledge of shoot branching at different levels of organization can theoretically be used to create a comprehensive mechanistic network model (Beveridge et al., 2023). However, this knowledge is derived from experiments conducted under different conditions and across various species, introducing significant noise and challenges for quantitative comparisons across experiments. To effectively leverage this noisy dataset from diverse experiments and species, an alternative modelling approach is necessary, one that differs from conventional methods designed around data generated specifically for the purpose of model development. Nevertheless, diverse experiments hold much information on qualitative relationships that are used to describe network topology.

Santolini & Barabási, (2018) demonstrated that network topology alone captures a substantial portion of perturbation dynamics without requiring kinetic parameters. Across 87 biological models, their DYNAMO framework showed that directionality was the critical topological feature, with directed networks achieving 65% accuracy compared to only 40% for undirected ones. Adding edge sign information distinguishing activating from inhibiting interactions provided only marginal improvement for predicting perturbation strength, but proved essential for predicting perturbation direction, where signed directed models reached 78% accuracy. For sparse, modular networks containing multiple strongly connected components, accuracy increased to approximately 80%. Maizels & Briscoe (2026) argues that networks assembled from statistical association collapse into uninterpretable “hairballs” that fail to capture causation. They set out three requirements for GRN models: mechanistic structure, coarse enough abstraction that not every molecular detail must be captured, and experimental perturbation as the basis for training and testing.

Building on this insight that topology encodes fundamental regulatory logic, Fortuna *et al*. (2026) developed a practical framework for translating network structure into testable mathematical models. The PSoup package, implemented in R, automatically converts signed directed networks into difference equations by applying a standardized set of algebraic rules that describe how nodes regulate one another from a mechanistic perspective. This translation from topology to mathematics creates a direct opportunity to simulate perturbations reported in the literature and compare predicted outcomes against observed experimental results. The algebraic rules underlying PSoup are grounded in ODE-inspired logic, as rigorously analyzed by Lawson et al., (2026), who used a pea branching network (Dun et al., 2009) to demonstrate how biological hypotheses about activation, inhibition, and synthesis at a homeostatic state can be systematically expressed as compact mathematical descriptions whose qualitative predictions are overwhelmingly determined by network structure rather than kinetic parameter values. PSoup places activating inputs in the numerator and inhibitory inputs in the denominator of each node’s equation to ensure a homeostatic baseline condition that means that the network remains stable regardless of the addition or removal of nodes. This enables qualitative predictions of network perturbations relative to a stable baseline and also enables scaling up. Genotypic switch variables such as knockout or overexpression control gene function. This design enables the automated construction of testable mathematical models directly from network diagrams (knowledge graphs), eliminating the need for manual equation formulation or kinetic parameterization and making perturbation testing accessible at the scale required for systematic literature-based validation.

Using PSoup, we constructed an evidence based highly curated topology of the regulatory network controlling shoot branching from prior biological knowledge and translated this topology into algebraic formulas to test whether it could predict perturbation outcomes reported in the literature. Because the quantitative outcome of shoot branch number is not suitable for direct comparison across experiments conducted under diverse conditions (Peters *et al*., 2016), we focused exclusively on the directionality of phenotypic change rather than its magnitude, reinterpreting quantitative data into qualitative terms. This approach is feasible because all experiments contain observations, such as the effects of mutations or treatments, that can be interpreted as qualitative changes relative to a wild type or control plant under control conditions. In this way, both experimentally and in the model, phenotypic differences emerge as effects of the network caused by differences among genotypes and treatments.

The objectives of this study are twofold. First, to assess the extent to which a directed signed network, constructed from prior knowledge in literature and translated into simple algebraic formulas, can predict the direction of change in the branching phenotype relative to the wild type at a homeostatic baseline. Second, to evaluate the predictive accuracy of the final network. To do this evaluation of prediction accuracy, we used independent data not involved in the model’s construction, including outcomes for nodes other than shoot branching itself. Together, these objectives test a central question: whether a well-curated causal topology in the PSoup algebraic construction (Fortuna *et al*., 2026), captures the essential regulatory dynamics of the biological system sufficiently to recapitulate phenotypic outcomes known across broad prior knowledge.

To achieve this, we constructed a theoretical shoot branching directed signed network based on prior knowledge from a literature search. We employed the PSoup modelling package (Fortuna *et al*., 2026) to translate the network into algebraic formulas following rules designed to compare perturbation outcomes relative to wild type. By simulating experimental treatments reported in the literature as perturbations within the network, we refined the theoretical network to achieve the best fit between simulated and observed biological outcomes regarding the direction of shoot branching change. We then assessed the predictive accuracy of this refined network by comparing simulated outcomes to biological results from experimental treatments not used during refinement, and for outcomes different from those considered during the refinement stage. Discrepancies between simulations and experimental observations are discussed for both the refinement and testing stages, along with how the approach exposed gaps in the prior knowledge. Finally, we discuss the potential use of these regulatory networks for crop improvement and fundamental discoveries.

## Materials and Methods

The shoot branching network model was developed through a systematic process summarized in Fig. 1. The process began with a comprehensive literature review to gather information on shoot branching regulation. This generated the training data set and enabled construction of an initial signed directed network based on prior biological knowledge. This network was then translated into mathematical equations using PSoup’s algebraic rules. The directed signed network was iteratively refined through manual curation until it achieved the highest accuracy in predicting the qualitative direction of shoot branching responses to perturbations reported in the literature, relative to baseline. Finally, the model was tested for its ability to predict directional changes in other network nodes beyond shoot branching, using independent perturbation data (testing data) not included in the training process.

**Fig. 1.**
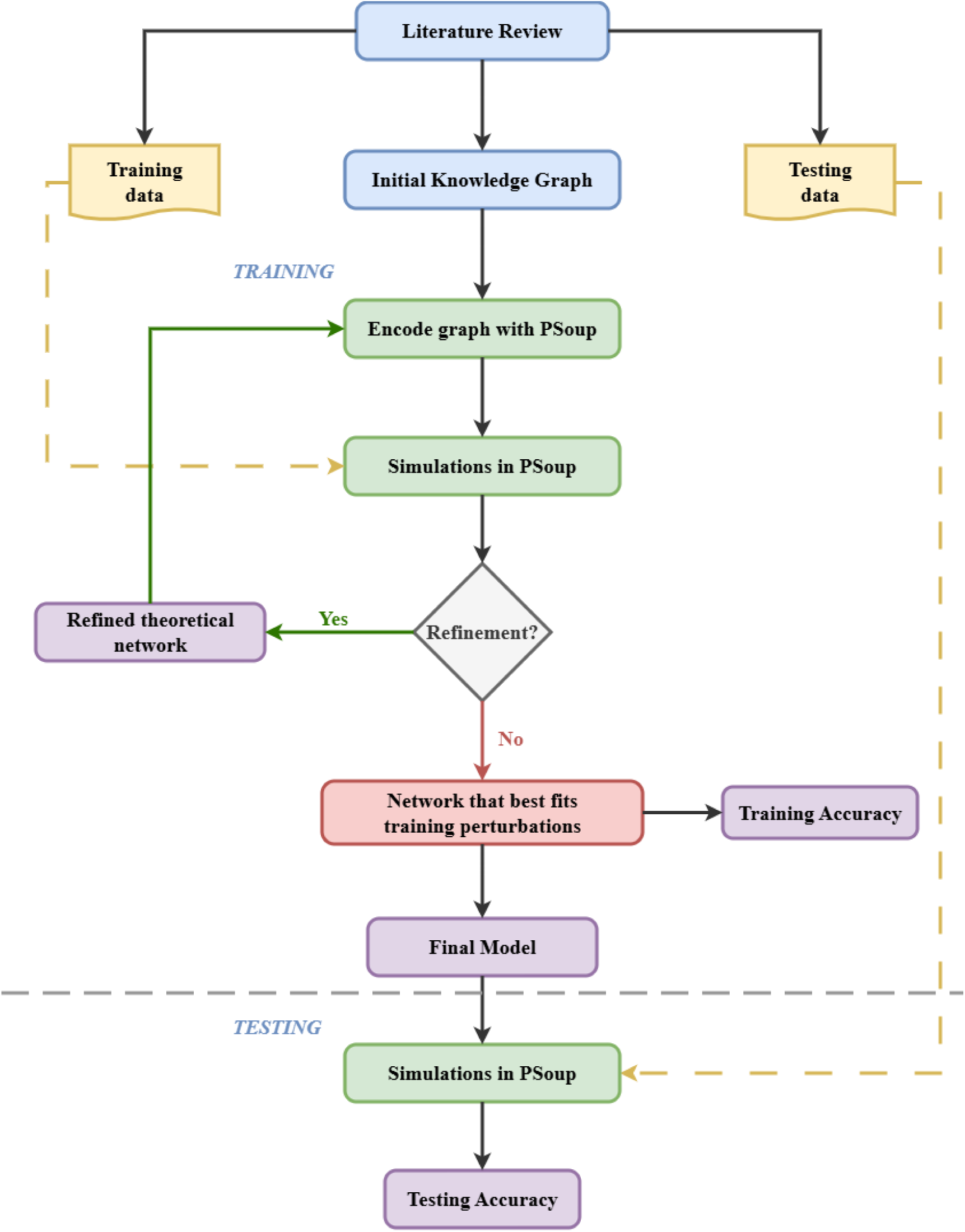
Flowchart of the model training and testing process. The process begins with a comprehensive literature review, which produces two outputs: (1) a knowledge graph constructed in Newt Editor, representing regulatory relationships between genes, hormones, and physiological processes as a signed directed network; and (2) biological perturbation datasets recording the phenotypic effects of mutants and treatments for use as training and testing data. The knowledge graph is exported in SBGN format and encoded into algebraic formulas using PSoup. Simulations replicate biological perturbations and the network is iteratively refined against the training data until accuracy is maximized. The final network model is then assessed on an independent testing dataset to evaluate its predictive accuracy.

### Literature review

A literature review was conducted to perform a domain analysis on branching in herbaceous plants. Relevant papers were identified through search queries in the Scopus database for the period from 2002 to February 2024, using terms related to shoot branching, tillering, and bud outgrowth in combination with mutant, knockout, overexpression, and hormone treatment terms (Supplementary Materials, Fig. S1, Table S1). Additional articles were included using a forward and backward snowballing method (Wohlin, 2014). Papers were included if they reported branching, tillering, or bud outgrowth phenotypes in herbaceous species with mutant or treatment comparisons to wild type. Studies conducted in woody species, review papers without original experimental data, papers lacking mutant or treatment comparisons to wild type, and those with insufficient quantitative data for qualitative interpretation were excluded (Supplementary Materials, Fig. S1, Tables S2 and S3). From an initial pool of 786 records identified through database searching and snowballing, 140 papers met the eligibility criteria (Table S2) for constructing the knowledge graph network (Supplementary Materials, Dataset S1). Of these, 49 reported perturbation data at the whole-plant branching phenotype level and were suitable for training (Supplementary Materials, Dataset S2), while 21 reported bud release and gene expression data and were reserved for testing (Supplementary Materials, Fig. S1; Supplementary Materials, Dataset S3).

Because the various publications used for extracting, training and testing perturbations spanned multiple herbaceous species, all gene and component names were manually translated into their Arabidopsis equivalents to ensure consistency across the network and perturbation datasets. This harmonization was necessary because gene names, pathway designations, and phenotypic descriptors vary considerably across Arabidopsis, rice, sorghum, and other species, with different research groups often using distinct terminology for orthologous genes and equivalent processes. All node names in the final network therefore use Arabidopsis nomenclature, with homologous and orthologous synonyms. Node names in the final network do not therefore infer that the data used to define the node has come from experiments in Arabidopsis.

The regulatory relationships between molecular and physiological components were identified from all 140 papers (Supplementary Materials, Dataset S1) and compiled into a knowledge graph using the Newt Editor platform. Nodes were defined as genes, hormones, metabolites, signalling proteins, or physiological processes with documented roles in branching regulation, while edges represented experimentally supported activation or inhibition relationships between these components, following the conventions of Fortuna *et al*. (2026). From the 49 training and 21 testing publications that contained perturbation data, phenotypic outcomes from genetic knockouts, overexpression lines, and exogenous hormone or sugar treatments were compiled to generate the training (whole-plant branching) and testing (individual bud lengths and gene expression) datasets, respectively (Supplementary Materials, Datasets S2 and S3). Each perturbation was recorded with its genotype, treatment conditions, and the observed qualitative direction of branching change relative to wild type.

### Encode network with PSoup

The directed signed network focussing on the whole-plant branching phenotype was constructed in Newt editor and exported in the Systems Biology Graphical Notation (SBGN) format (Le Novère *et al.,* 2009) as per Fortuna *et al*. (2026). This network was then converted into algebraic formulas using PSoup (Fortuna *et al*., 2026). The conversion of the networks to algebraic formulas followed algebraic rules designed to capture the network’s responses to perturbations from the baseline wild type (WT) condition. For further details and tutorials on PSoup, refer to the official documentation https://nicolezfortuna.github.io/PSoup and Fortuna *etal*. (2026).

### Model Equations

The network equations define the steady-state update rules for each of its 32 nodes. PSoup automatically translates the signed directed network topology into algebraic equations following a general form, where stimulatory inputs appear in the numerator and inhibitory inputs appear in the denominator. Under WT conditions, where no perturbations are present, the network remains in a steady state, with all elements assigned a value of 1. After perturbation of the network, the system iterates until all node values converge to a steady state. At each iteration *t*, the value of every node is computed from its regulators at iteration *t*− 1.

For each node X, the genotype modifier *g*with subscript *X*controls the genetic state of the node. Under wild-type and/or control conditions `g` equals 1 for all nodes. A knockout is represented by setting `g` to 0, and overexpression or exogenous supply by setting *g* greater than 1. Nodes with multiple stimulators or inhibitors use the arithmetic mean of their respective inputs. Some nodes require a necessary input, which is multiplied directly into the equation so that loss of the necessary input drives the node to zero regardless of other regulators. The general PSoup equation form is:

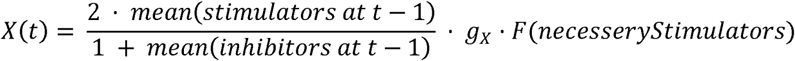

where *mean(stimulators)* is the arithmetic mean of all stimulatory inputs at the previous iteration, *mean(inhibitors)* is the arithmetic mean of all inhibitory inputs at the previous iteration, and *F(necessarystimulators)* is the function used by PSoup to simulate necessary stimulators. This formula ensures that when all inputs and the genotype modifier are at their wild-type value of 1, every node is equal to 1. The factor of 2 in the numerator compensates for the 1 in the denominator, maintaining the wild-type steady state. The 1 in the denominator also prevents problems when the inhibitor takes a value of 0. The *Shoot Branching*network (Fig. 2) translates into the equations below through PSoup. These equations are presented as raw PSoup output without compaction, for transparency.

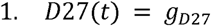

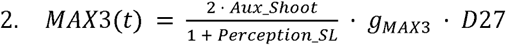

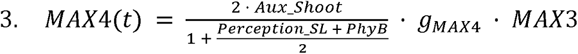

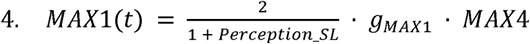

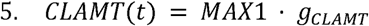

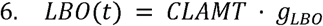

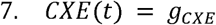

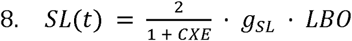

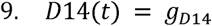

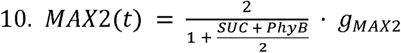

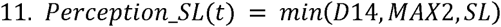

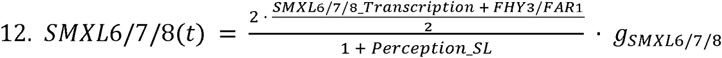

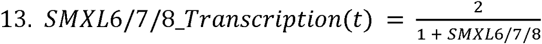

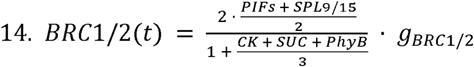

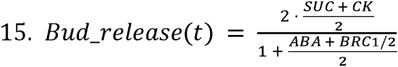

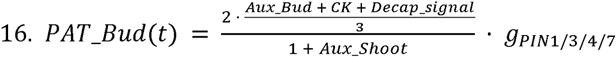

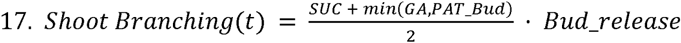

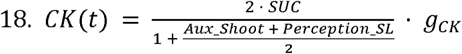

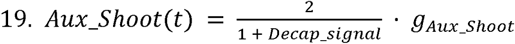

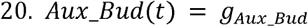

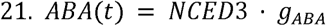

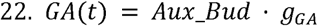

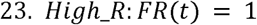

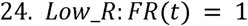

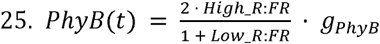

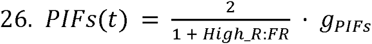

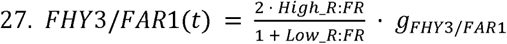

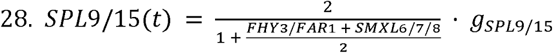

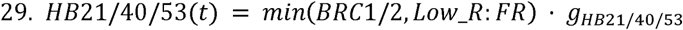

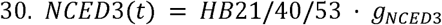

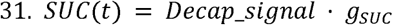

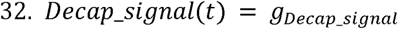

**Fig. 2.**
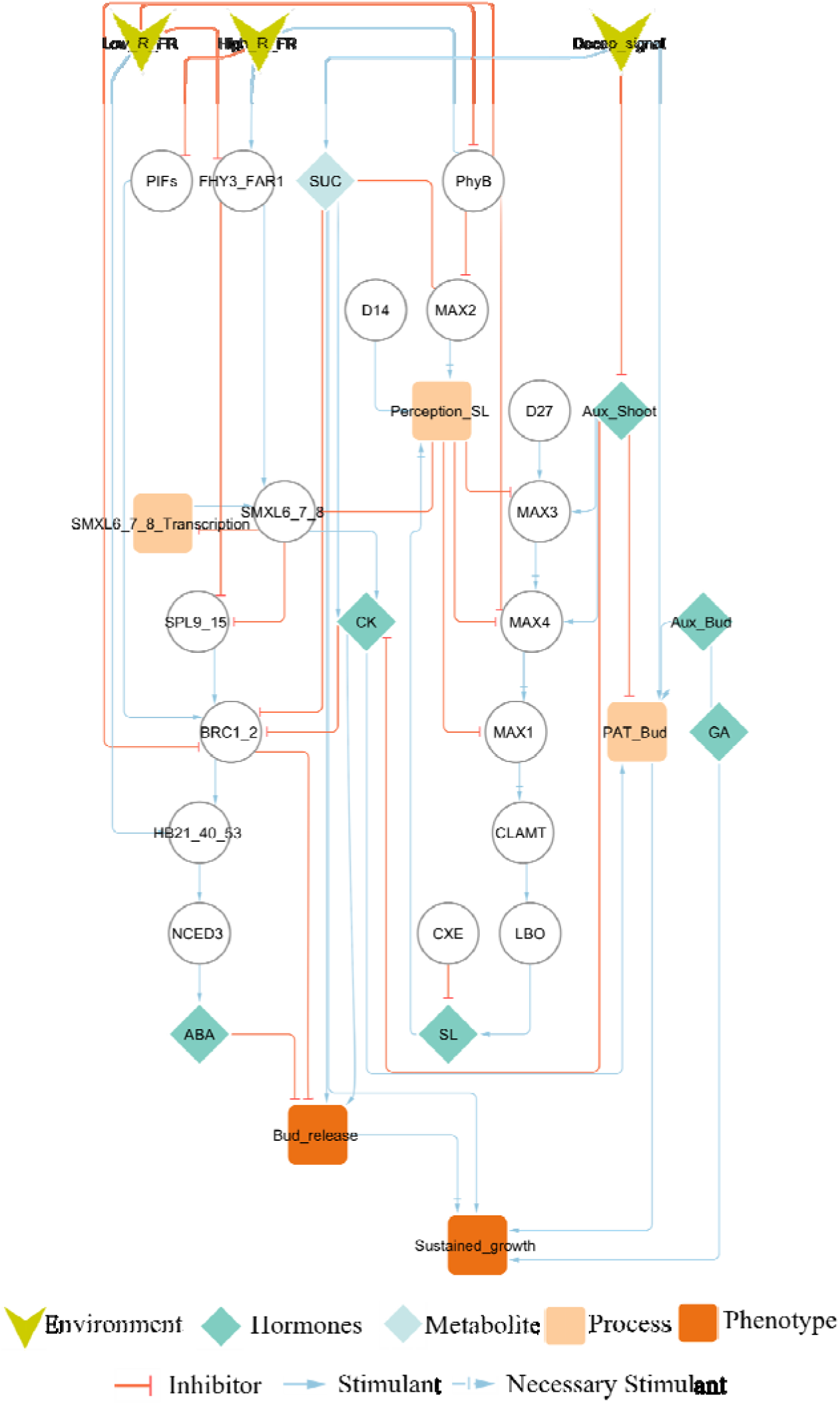
The final theoretical shoot branching network. Nodes represent genes, transcripts, hormones, and physiological processes such as polar auxin transport (PAT Bud), environment; edges represent regulatory interactions (activation or inhibition). Input nodes include decapitation and light quality. The network therefore integrates multiple organizational levels including hormone signalling (strigolactones, cytokinin, auxin, ABA, gibberellin), sugar supply, and light quality. Shoot branching was the output node for the training data set and was a measure of whole plant branching. Bud release which was focused on early branch growth typically at a single node as well as gene expression at multiple nodes of the network were used for the testing data set. The network in this figure is depicted using cytoscape for its enhanced clarity; necessary stimulants are highlighted with cross delta symbol. For the original figure in SBGN form (See Supplementary Materials, Dataset S18).

### Simulations and training

Simulations in PSoup were performed as perturbations designed based on biological perturbations reported in the literature. These biological perturbations, such as mutants or mutants combined with exogenous hormone treatments, were simulated relative to controls or wild-type, and the simulation results were compared against the corresponding experimental outcomes. The training dataset comprised 304 perturbations, of which 84 were unique (49 papers; Supplementary Materials, Dataset S2). Of the 84 unique types of perturbations to the network, four showed inconsistent results within or across species or lacked replication (Supplementary Materials, Table S4; Supplementary Materials, Dataset S4) and were therefore excluded, leaving 80 perturbations with biological consensus; a further two perturbations were excluded as erroneous records introduced during export, resulting in 78 unique perturbations used for evaluation. The training data was predominantly derived from experiments conducted on Arabidopsis and rice, with a few cases from petunia, pea, switchgrass and maize with a focus on data on the whole plant branching phenotype.

The model predicts whether the shoot branching phenotype increases, decreases, or remains unchanged in comparison to the WT or control after perturbations. The outcome of simulations was therefore tested by comparing the direction of change at the *Shoot Branching* node with the direction of change in shoot branching in the training data (Fig. 2). The initial model and corresponding knowledge graph were manually refined over iterative cycles until it most accurately reflected the biological perturbations (Fig. 2).

### Model accuracy

Model accuracy was determined by comparing the predicted direction of *shoot branching* change with the experimentally observed biological outcome for each perturbation. Each comparison was classified into one of three categories using a simple bin method as described below. A prediction was considered correct when the predicted bin matched the biological bin. It is important to note that not all perturbations were compared to WT. Instead, the baseline chosen for each comparison followed the experimental design of the source publication (Supplementary Materials, Supplementary Text 1).

Bin method to express direction of the simulation

The bin method is a statistical sign test that evaluates whether the outcome of a perturbation is higher, lower, or the same as the WT condition, and whether it aligns with the biological perturbation data. This approach was previously utilized by Dun et al., (2009) and exemplified again in Lawson *et al.,* (2026) and Fortuna *et al*. (2026).

Specifically, three bins were defined:

- −1: The outcome of the output node was lower than the control.
- 0: The simulated value was within 5% of the control (ratio between 0.95 and 1.05).
- +1: The outcome was higher than the control.

This method tests whether a perturbation results in an increased, decreased, or unchanged phenotype (branching for training; bud length or gene expression for testing) relative to the control. Because the model is qualitative and predicts the direction of change rather than its magnitude, a prediction was scored as correct when it matched the biologically reported direction: for a reported increase or decrease, when the simulated value moved in the same direction; for a reported no change, when it stayed within the 5% no-change band. A 5% band, rather than exact equality, was used because in a signed network where stimulatory and inhibitory paths converge on a node the opposing contributions almost never cancel exactly, so one path is usually marginally stronger and leaves a small residual deviation that does not represent a genuine change.

### Threshold selection

The binning threshold determines the PSoup simulation output range around the control value within which a perturbation is classified as unchanged. When a perturbation is simulated, the model produces a numerical value under both perturbed and wild-type/control baseline conditions, and the ratio of these values determines the direction of change. If the perturbed value exceeds the wild-type value by more than the threshold, *shoot branching* is classified as increased, and if it falls below by more than the threshold, it is classified as decreased, with all remaining cases classified as unchanged. Low thresholds will produce over-sensitive classifications by assigning directionality to minor numerical fluctuations, while high thresholds will miss classify genuine changes by classifying them as unchanged (Supplementary Materials, Fig. S2).

### Model testing against an alternate dataset

In contrast to the training dataset, which focused on the number of branches per plant, the testing dataset focused on bud length at a given node and gene expression data. It consisted of 135 perturbations, of which 88 were unique and 84 of these unique perturbations had biological consensus and were used for testing (Supplementary Materials, Datasets S3 and S5), enabling an independent evaluation of the model’s predictive accuracy. The testing data compared PSoup predictions against biological consensus derived from multiple experimental replicates (Supplementary Materials, Dataset S3).

### Randomised rewiring of the curated topology

To test whether the topology of the curated network matters for its testing accuracy, and whether that accuracy could be reproduced by a network whose wiring of the topology had been randomised, we simulated the same perturbations in randomised networks and tested them in the same way as the shoot branching network. This approach has been used for assessing biological networks and involves randomizing one or more aspects of wiring within a given topology (Maslov & Sneppen, 2002; Ortiz-Gutiérrez *et al.,* 2015). By wiring within topology we mean the wiring within the network structure, which includes its edges, its sign and its influence type. We therefore did not compare the model against arbitrary random networks but against similar networks. We used four randomisation schemes, ranging from complete randomisation to conservative rewiring of a few edges, each retaining the same 32 nodes and regulatory logic and differing only in wiring (Supplementary Text 4).

We used four randomisation schemes, each retaining the same 32 nodes and the same regulatory logic and differing only in wiring, ordered here from the scheme that preserves most of the curated structure to the scheme that preserves least. Sign shuffling leaves every connection including the directionality between the same two nodes and permutes only whether the connection activates or inhibits. Signed-degree-preserving rewiring moves the connections but keeps, for each node, its number of incoming and outgoing connections. Degree-preserving rewiring keeps only the number of incoming and outgoing connections per node. Erdős–Rényi rewiring keeps only the nodes and the number of connections of each influence type and joins them at random. We performed these randomisations to test whether each component of network topology, the edges, their direction and their sign, contributes to predictive accuracy. Each scheme was run 1000 times (Supplementary Text 4).

## Results and Discussion

### Model development

Following the flow outlined in *Error! Reference source not found.*, we created an initial network based on a literature review (Supplementary Materials, Fig. S1) including the most recent published reviews (Beveridge et al., 2023; Dun et al., 2023) and focused down on a core group of publications used to create a theoretical branching network of the type in Fig. 2. We then used PSoup to translate the network into mathematical formulas (see methods Model Equations). The model was tested on whether it could predict the relative branching response of each perturbation against its own baseline, which differed across perturbations. (Supplementary Text 1). The dataset includes mutant treatments, mutants combined with exogenous hormone supply, and wild-type plants supplied with exogenous hormones or metabolites.

### Model training

The network was iteratively refined through 89 manual curation cycles, testing alternative configurations until the model achieved optimal accuracy in fitting perturbation outcomes. The final network (Fig. 2) achieved the highest accuracy in predicting shoot branching outcomes, and further modifications did not lead to improvement. This indicates that the model represents an optimal configuration for this dataset while maintaining the requirement that all edges have optimal agreement with the published literature (e.g., reviewed in Beveridge *et al*., 2023; Dun *et al*., 2023). The final network (Fig. 2) achieved 86% accuracy in predicting the observed direction of shoot branching change relative to baseline (Fig. 3).

**Fig. 3.**
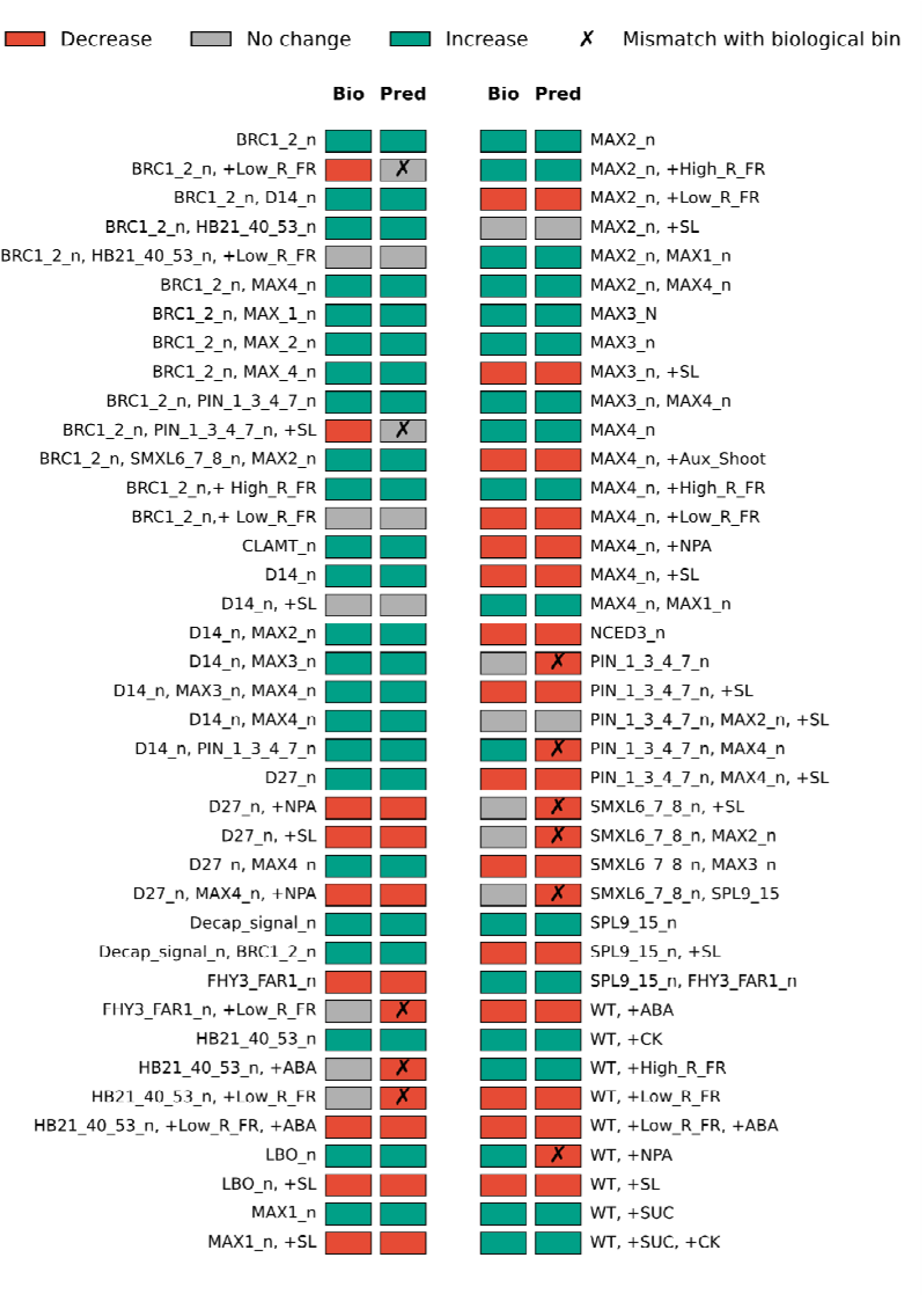
Perturbation comparison of the expected biological direction against the PSoup-predicted direction for the 78 unique training perturbations from Dataset S9. Each row shows one perturbation with two coloured squares, where the left square (Bio) is the literature-curated biological bin and the right square (Pred) is the PSoup-predicted bin based on the direction of change of perturbation relative to control ShootBranching(5% no-change band). Decrease (−1) is shown in red, no change (0) in grey and increase (+1) in teal. A black cross on the predicted square marks the rows where the PSoup prediction does not match the biological bin. Mutants are represented as perturbations to nodes written as GENE_n. The model agrees with the biological bin in 67 of 78 perturbations (86%). The modifier and exogenous values and the raw results for the simulations are in (Dataset S6, S7 and S8).

### Per-class accuracy

We used a confusion matrix to show how well the model performs by comparing actual vs predicted classes. The confusion matrix and per-class metrics (Supplementary Materials, Table S5) show that the model performed best for class +1 (increase; F1 = 0.98), followed by class −1 (decrease; F1 = 0.80). Class 0 (unchanged) had the lowest F1 score (0.53), indicating that the model tends to better predict changes rather than no change. This is expected given the qualitative nature of the approach: perturbations near the threshold boundary are inherently difficult to classify, as small simulation values close to the control value may fall on either side of the threshold. The lower F1 score for class 0 (unchanged) is also expected for a more fundamental reason. Statistical tests can demonstrate that two conditions are significantly different, but they cannot confirm that two conditions are truly the same. When a biological experiment reports no significant change, this may simply mean the difference was not detectable given the number of available replicates, measurement precision, and/or the nature of the assay. Some experimental observations classified as unchanged may therefore involve genuine biological changes that the experiment lacked the power to detect. When the model predicts a directional change for such cases, it is penalized as an error even though its prediction may be biologically correct, and the lower F1 for class 0 should be interpreted with this inherent limitation of the benchmarking data in mind.

### Threshold sensitivity

The binning threshold determines the range around the control value within which a perturbation is classified as “unchanged.” We evaluated a range of thresholds from 0.005 to 0.200 and computed accuracy at each threshold. A threshold of 0.05 was selected, representing a 5% deviation from baseline, as it gave a high overall accuracy of 86 % for this specific network and dataset on a stable performance plateau, though the optimal threshold may differ for other networks (Supplementary Materials, Fig. S2).

### Discrepancies due to ambiguous pathways in the shoot branching network

Of the 78 unique training perturbations, the model reproduced the experimental direction of shoot branching in all but eleven (Fig. 3; Supplementary Table S6). Of these eleven, four involved auxin transport (two via the PIN proteins, one from NPA treatment of wild type, and one a *brc1 pin1/3/4/7* double treated with SL), three concerned R:FR light signalling, one was linked to ABA pathway components and three arose from *smxl6/7/8* perturbations. Three of these eleven are multi-mutants, *pin1/3/4/7 max4, smxl6/7/8 max2* and *smxl6/7/8 spl9/15*, which we compared with the wild type for consistency. Biologically this high throughput comparison against WT is not the appropriate contrast, since a higher order mutant should be compared not with the wild type but with the relevant lower order mutant, for example a double against its corresponding single. Under this comparison the model predicted the correct direction for three of the four multi-mutant comparisons (Supplementary Materials, Table S8) with the model then failing only for for *smxl6/7/8 spl9/15* against *spl9/15*. These corrected mismatches therefore reflected the choice of reference genotype (WT) rather than a failure of the model. The reason for the remaining eight mismatches out of the original 78 perturbations in the training data set are discussed below.

### Inaccuracies due to branching values being close to WT

Two discrepancies were in the scenario where the model predicted a decrease but the experimental difference was not significantly different from WT and in both cases it was due to the control already having very few branches. For the *pin3/4/7* mutant, the model predicted decreased *shoot branching* relative to wild type Arabidopsis, whereas the experiment showed no change (Fig.3B in van Rongen et al., 2019). In these cases, wild type *shoot branching*is already low, and a further decrease may be difficult to capture as a significant difference due to the branching number trait being measured as whole integers. The model predicted decreased branching in *fhy3 far1* mutant plants under low R:FR compared to standard light conditions while the experiment reported no change (Fig. 1b in Xie et al., 2020). *fhy3 far1* plants already have very few branches in white light (∼1 per plant), so shade cannot reduce them further, or not with a significant difference.

### R:FR light signalling

Two discrepancies involved low R:FR treatments applied to *brc1* (; Table S6). In *brc1* under low R:FR the model predicted no change relative to *brc1*, while the experiment showed that branching is still reduced when *brc1* plants are moved from white light to white light supplemented with far-red (Fig.3B in González-Grandío et al., 2017). This is because *BRC1* node has already been removed in the *brc1* mutant, the low R:FR signal has lost its only route to branching, because the effects of low R:FR on bud activity are channelled through *BRC1*. With this node knocked out, supplementary FR has no remaining path by which to influence the phenotype, and so the model predicts that *brc1* and *brc1* under low R:FR are the same. This indicates an area for further experimentation.

The second R:FR discrepancy involved high R:FR applied to the *brc1* mutant. In this experiment planting density was the treatment tested, and to represent it we simulated low density as a high R:FR ratio, since the ratio rises as the spacing between plants increases. The model predicted an increase in branching relative to the *brc1* control, consistent with the direction reported experimentally (Figure 7A in Aguilar-Martínez *etal.,* 2007), but the predicted change fell within the 5% no change band and was therefore automatically binned as no change.

### ABA pathway

The single ABA-related discrepancy concerns the *hb21 hb40 hb53* triple mutant treated with exogenous ABA, compared to the triple alone. The experiment reported no change in branching (Fig. 4E in González-Grandío et al., 2017), while the model predicted a decrease (Supplementary Materials, Table S6). In the network, *HB21*, *HB40*, and *HB53* promote ABA, which inhibits bud outgrowth, so knocking them out removes this inhibition and the model branches more. When ABA is re-added in the knockout, the ABA node reactivates and the model restores the inhibition, lowering branching. The experiment instead shows no change, meaning the *HB* genes are required for ABA to act on the bud rather than only for its production. The model is not incorrect here, since it represents the topology faithfully and the ABA node does activate when ABA is re-added. The limitation is that it treats the ABA-to-outgrowth effect as fixed rather than conditional on the *HB* genes. This can be captured by encoding ABA’s effect on outgrowth as an AND relationship that requires both ABA and the *HB* genes, placing the *HB* genes as a co-requirement at the outgrowth step in addition to their upstream role in producing ABA. When we implemented it this way, it corrected this case but reduced the accuracy of others that were already correct, so we did not keep it, to avoid overfitting a relationship that the network otherwise models correctly. Consequently, we recommend that further experimentation is needed.

**Fig. 4.**
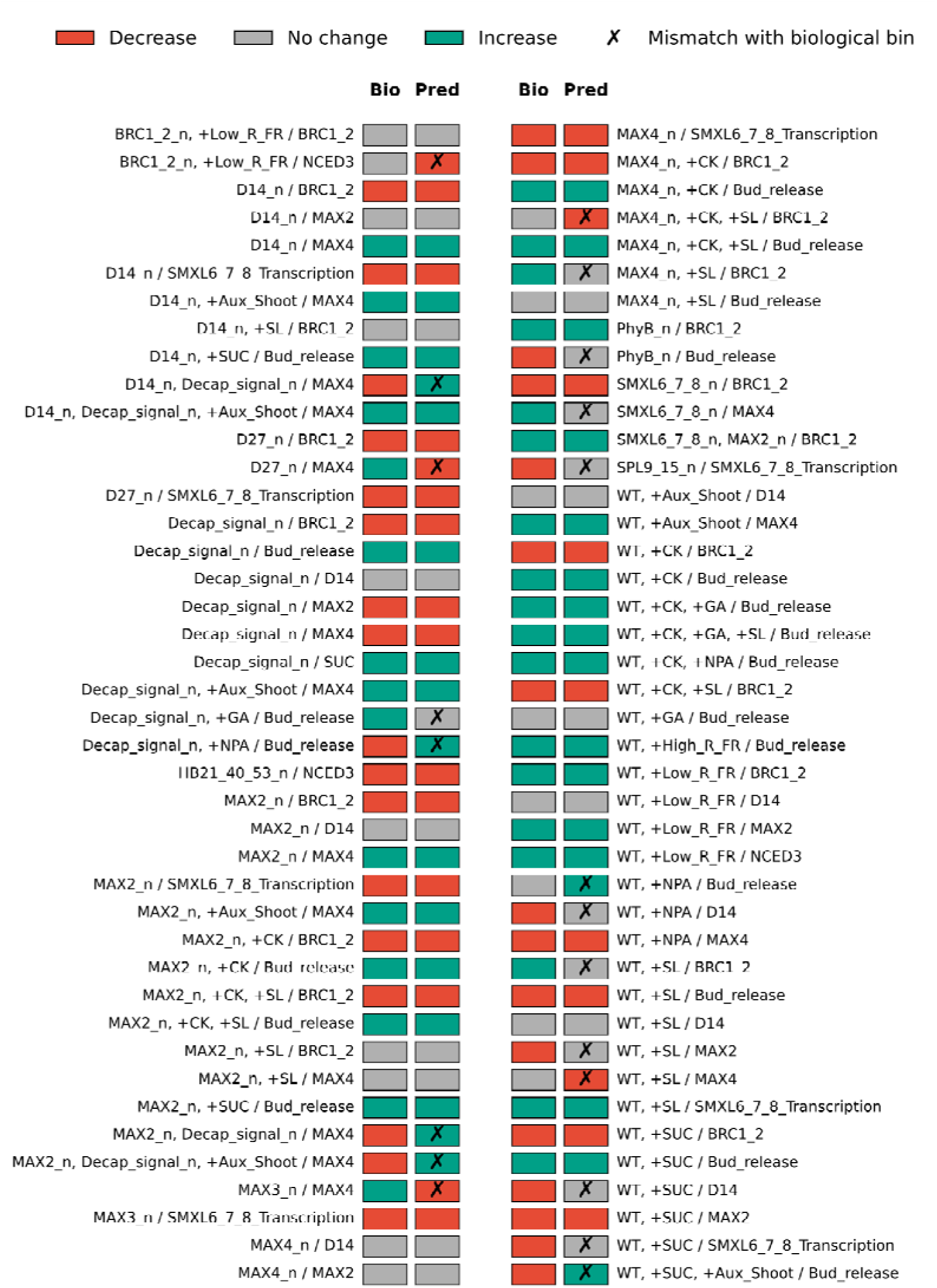
Per-perturbation comparison of the expected biological direction against the PSoup-predicted direction for the 84 unique, biologically consistent perturbation-node pairs in the testing set from Dataset S13. Each row shows one (perturbation / measured node) combination with two coloured squares, where the left square (Bio) is the literature-curated biological bin and the right square (Pred) is the PSoup-predicted bin at the measured node, based on the direction of change (5% no-change band). In this figure, perturbations in genes indication mutations while a measured node labelled with a gene refers to the gene expression. Decrease (−1) is shown in red, no change (0) in grey and increase (+1) in teal. A black cross on the predicted square marks the rows where the PSoup prediction does not match the biological bin. The model agrees with the biology in 63 of 84 pairs (75%). The modifier and exogenous values and the raw results for the simulations are in (Dataset S10, S11 and S12).

### Auxin Transport

This same single-node abstraction of polar auxin transport also underlies the one training mismatch introduced by representing NPA as both a block of PIN-mediated transport and a reduction in shoot auxin: wild type treated with NPA branches more in the literature (Lin *et al*., 2009), but because knocking out the *PAT_Bud* node prevents bud auxin export the model instead predicts less branching. Capturing this opposing outcome would require the spatial, canalisation-based representation of auxin dynamics (Prusinkiewicz *et al*., 2009; Nahas *et al*., 2025) rather than the single lumped transport node used here.

### SMXL6/7/8 with SPL9/15 and D53

For smxl6/7/8 spl9/15 the model predicted decreased branching despite an unchanged phenotype relative to wild type (Fig. 6b in Xie et al., 2020). The lower order comparison (against spl9/15) gave the wrong direction of change because the perturbation was represented in a linear pathway of shade - FHY3/FRY1 - SMXL6/7/8 - SPL9/15 - BRC1 - branching. The model had not implemented the parallel route of shade - PIF - MIR156 - SPL9/15 (Xie et al., 2020) and doing so would require additional modelling and testing.

Due to the extremely low branching phenotype of smxl6/7/8mutants it is not informative to add exogenous SL in order to inhibit branching further. However, the D53 mutant in rice is a constitutive over expression line of this node and is unable to be degraded by SL and hence SL has no inhibitory effect on branching in D53 mutants (Fig. 1E in Song et al., 2017). Our model is built for SMXL, so this scenario does not map cleanly onto how the D53 perturbation should be represented.

### Epistasis analysis shows the gap of parallel pathways in literature in different pathways

The standard training analysis compares all perturbations to wild-type or baseline controls, giving 86% accuracy (67/78) overall and 86% (19/22) on the multi-mutant subset (treatments carrying two or more gene mutations, such as double and triple mutants). However, this can mask incorrect predictions for multi-mutants, because comparing a multiple mutant directly to wild type does not isolate the effect of the added mutation on top of the single-mutant background. This was discussed above for *pin1/3/4/7max4, smxl6/7/8max2*where the correct comparison to the appropriate single mutant improved the model accuracy. Therefore, we sought to use this approach across the whole training data set. Under an epistasis-corrected analysis, in which each multi-mutant is compared to its relevant lower-order mutant control rather than to wild type, accuracy on this subset dropped to 59.1% (13/22; Supplementary Materials, Table S8). In nine of these comparisons, the wild-type baseline had produced correct predictions, but only because it did not account for epistatic interactions between the combined mutations (Supplementary Materials, Supplementary Text 2; Tables S8; Datasets S16 and S17). Importantly, seven of these newly exposed disagreements involved double knockouts within the SL biosynthesis chain. The other two were double mutants, involving *PIN*and *SPL9/15* genes, which the model failed to predict correctly in the training set as well. With wild-type baseline, double mutants such as strigolactone deficient *max3 max4* appeared correctly predicted because both the single and double mutant show increased shoot branching relative to wild type. However, when compared to the single mutant (e.g., does *max3 max4* branch more than *max3* alone?), the model incorrectly predicts no additional effect, whereas experiments show enhanced branching. This result supports the hypothesis that MAX3 and MAX4 produce non-canonical metabolic products with SL-independent branching activity (Bruno *et al*., 2014). This aspect of SL biosynthesis is commonly represented in diagrammatic representations of the branching network (Bennett *et al*., 2016; Beveridge *et al*., 2023; Domagalska & Leyser, 2011; Dun *et al*., 2023; Rameau *et al*., 2015; Wang *et al*., 2020). The *max3* and *max4* mutant alleles used in these studies are T-DNA or transposon insertion mutants generally considered to be complete loss-of-function, so the additive double-mutant phenotype is unlikely to reflect residual enzymatic activity and instead points to independent pathway contributions. Importantly, while the original studies reporting these double-mutant phenotypes captured the additive effects experimentally, their primary objective was not to identify the specific parallel pathways responsible for the enhanced phenotype. The modelling approach presented here provides an additional framework for exposing such gaps, by formalizing the known linear pathway and testing it against experimental data. The model pinpoints exactly where the linear assumption fails, thereby generating testable hypotheses about unmapped parallel biosynthetic routes. This capacity to identify missing pathway components represents a key advantage of qualitative network modelling for guiding future experimental work.

Additional parts of the network revealed by this approach included double mutants between SL mutants and *brc1*. Aguilar-Martinez *et al*. (2007) reported that these double mutants actually had slightly fewer branches than one or more of the single mutants, which is not consistent with the canonical branching networks (review papers) or our model (Table S8). Seale *et al*. (2017) produced combined SL mutants with *brc1 brc2* double mutants and this multi-mutant was indeed more highly branched than the SL mutants or *brc1 brc2*. This is consistent with the model output.

The final remaining multi-mutant that produced an incorrect prediction was for the *brc1 pin1/3/4/7* mutant (Table S8). The model predicted that the multi-mutant would have less branches than *brc1* yet van Rongen *et al*. (2019) reported that there is no difference in branching phenotype between these lines. They showed that whereas *pin1/3/4/7*represses branching in *max2* and *max4*, it cannot do so in *brc1 brc2*. This highlights an area worthy of more attention (Nahas *et al*., 2025; van Rongen *et al*., 2019).

### Summary of accuracy against the training model (Shoot branching)

The accuracy of the model, tested against a baseline of WT for mutations, or WT or the mutant background for exogenous treatments (for example a double mutant plus an exogenous treatment compared with the double mutant alone), was 86% - but see the caveats on interpreting this percentage as discussed below. This comparison provides a rapid and convenient measure of accuracy and lets us see the emergent properties of the network as a whole, but it is limited for cases where multiple combinations of gene perturbations are used. In those cases, the multi-perturbation treatments can instead be compared against perturbations with one fewer treatment, such as double mutants compared with single mutants. When doubles were compared with singles, the model correctly predicted the intermediate phenotypes of doubles acting across different pathways, but not those within the same linear pathway, such as the SL pathway. Comparing multi-perturbation treatments against those with one fewer treatment perturbation is therefore useful for testing the wiring of the network.

### Model testing

The final shoot branching network that scored the highest accuracy in training (Fig. 2) was fitted exclusively to predict the direction of shoot branching change, measured as the number of branches per plant. To test whether the network’s algebraic rules could generalize beyond this single phenotypic output, we evaluated the model’s ability to predict directional changes at other nodes throughout the network, including the *Bud release* phenotype and gene expression levels for *BRC1/2, MAX4, MAX2, D14, NCED3, and SMXL6/7/8* (Fig. 4). None of these node-level predictions were used during training, so there was no opportunity to adjust the model topology to fit these data.

*Bud release* refers to the early transition of axillary buds from dormancy to active growth. The measurement of *bud release* varies across studies, with some studies reporting the proportion of nodes along the stem with any measurable outgrowth compared with controls, and others measuring bud length beyond a defined minimum threshold such as a 5 mm. The evaluation set comprised 84 unique perturbation-node pairs within the model aggregated from 135 experimental observations across 21 papers and five species, predominantly Arabidopsis, pea, rice, rose and sorghum (Supplementary Materials, Datasets S3 and S5).

### Testing accuracy

The model matched the experimental direction for the testing data in 63 of 84 perturbation-node pairs, giving an overall testing accuracy of 75% (Fig. 4; Supplementary Materials, Table S7). Twenty-one predictions did not agree with the literature (Supplementary Materials, Dataset S13). We grouped the mismatches into categories describing the possible reasons that perturbations from prior knowledge could not be predicted.

### Spatial or temporal dynamics

Most of the discrepancies in the testing data set (15 of 21) involved treatments with exogenous synthetic hormones or sugars (Supplementary Materials, Dataset S7), most likely reflecting factors such as the timing, concentration or tissue specificity of gene expression measurements that fall outside the scope of the current qualitative model, which does not explicitly represent spatial or temporal dynamics. Some of these are also discussed in the following sections.

### Topology not implemented

In this group perturbations include mismatches due to missing or inaccurate topology such as such as feedback regulation of gene expression, or a mismatch between gene expression and protein or product levels. In ten cases the incorrect predictions were because the model trained on shoot branching data does not represent transcript levels independently of the enzymatic (biochemical) or gene product (protein) levels which are used to affect downstream nodes (Supplementary Materials, Table S7). This issue accounted for any mutants that affected nodes upstream of MAX4 (such as MAX3) or downstream, such as SMXL6/7/8 and treatments that modify SL levels or signalling including SL treatments or treatments that modify auxin content such as NPA. These treatments all affect levels of gene expression that do not then lead to changes in branching. For, example, low SL levels or signalling lead to feedback upregulation of several SL biosynthesis genes (Yao *et al*., 2026), even though this will have no effect on branching if the SL pathway is blocked through mutation. This information is not included in the training model which focuses on results impacting the total branch number.

The rice *d27* mutant, the *d17* mutant (MAX3 in the network) and the *d53* mutant (SMXL6/7/8 in the network) each reported increased *D10 (*MAX4 in the network) gene expression levels relative to WT whereas the model predicted decreased *MAX4* gene expression. As discussed above, this is mostly due to the fact that feedback regulation of the expression of strigolactone biosynthesis genes is not implemented in the model.

Feedback regulation of SL levels from SL perception is implemented in the model via single SL nodes that effectively represent only the enzymes, not the transcript levels. Therefore, SL biosynthesis gene expression predictions will not be accurate if this part of the network is also perturbed such as in treatments with the pea SL signalling mutants (affecting MAX2 and D14). These inaccuracies included these mutants combined with decapitation or auxin treatment (Table S7 verses Figure 7 in Foo *et al.,* 2005).

The remaining mismatches in topology (Supplementary Table S7) arose because this level of topological detail was not implemented in the model during training. Decapitated WT Arabidopsis seedlings treated with exogenous NPA reported decreased *D14* gene expression relative to decapitated WT (Figure 6C in Chevalier et al., 2014), whereas the model predicted no change. WT rice treated with exogenous sucrose reported decreased *D14* relative to WT (Fig. 3c in Patil et al., 2022) whereas the model predicted no change, because the shoot branching network implemented does not include a sucrose-to-D14 connection and therefore cannot reduce D14 in response to sucrose. WT rice treated with exogenous sucrose also reported decreased *D53* transcript relative to WT (Fig 3b in Patil et al., 2022), whereas the model predicted no change, because the SMXL6/7/8 transcription node in the network depends only on the SMXL protein state and has no edge from sucrose to represent the transcriptional reduction observed in rice. The rice *ipa1* (SPL9/15 in the network) mutant reported decreased *D53* transcript relative to WT (Figure 6E in Song *et al*., 2017) whereas the model predicted no change, because the network has no edge from SPL to SMXL6/7/8 transcription to represent the SPL-driven activation of *D53*. WT Arabidopsis treated with exogenous SL reported decreased *MAX2* relative to WT (Figure 6B in Chevalier *et al.,* 2014), whereas the model predicted no change, because the shoot branching network has no feedback from Perception_SL to MAX2 and therefore cannot represent the SL-induced degradation of MAX2. The Arabidopsis *brc1* mutant under low R:FR reported no change in *NCED3* transcript relative to WT whereas the model predicted decreased NCED3, because the network routes all NCED3 regulation through BRC1, so losing BRC1 also collapses NCED3 to zero in silico (Fig. 4A in González-Grandío *et al.,* 2017).

### Not Statistical significance

Mismatches in this group were due to the fact that the direction was correct but in the experiment the statistical difference was not enough to declare the difference. This issue is similar to that discussed earlier regarding the small inhibition of branching possible in WT plants. For the testing data set, in wild type Arabidopsis treated with exogenous SL, *MAX4* transcript was reported as unchanged because the decrease relative to wild type did not reach statistical significance (Fig 6B in Chevalier et al., 2014). The trend was nonetheless a decrease, which matches the reduction predicted by the model. The mismatch therefore reflects a non-significant experimental trend rather than a wrong directional call.

#### Comparison with intermediate phenotypes

In this group the mismatches arise because we compared each multi-perturbation treatment against wild type. When these treatments are instead compared against a perturbation with one fewer treatment, for example a double mutant against its corresponding single mutant, the intermediate phenotype is predicted correctly. The only cases that were incorrectly predicted after this improved comparison for the testing data set are discussed below.

For example, in WT rose treated with exogenous sucrose plus exogenous auxin (Fig. 2c in Bertheloot et al., 2020) reported decreased *bud release* relative to wild type, whereas the model predicted increased *bud release*. In the network, sucrose is the more dominant node and drives bud release upward, overriding the auxin input. The pea *rms1 (*MAX4 in the network) mutant treated with exogenous SL, and the pea *max4* mutant treated with exogenous cytokinin together with SL, reported increased and unchanged *BRC1* transcript relative to WT (Figure 5 in Dun *et al.,* 2012) whereas, relative to rms1, the model predicted unchanged and decreased respectively BRC1. However, again if the double treatment +CK and +SL are compared with the minus one treatment which is the +SL only then the model predicts the intermediate phenotype (Figure 5 in Dun *et al.,* 2012; Figure 1 in Kerr *et al.,* 2020; Figure 4B in Fang *et al.,* 2020).

### Below cut-off

In this group the mismatches arise because we defined a difference from baseline of 0.05 as the threshold for change, and any response falling below that cut-off is recorded as unchanged. In these cases, the direction of the simulated response can still be correct, for example an increase, but the magnitude is not large enough to cross the threshold and register as a difference.

WT Arabidopsis and pea treated with exogenous SL reported increased *BRC1* gene expression relative to WT (Figure 5 in Dun et al., 2012; Figure 1 in Kerr et al., 2020; Figure 4B in Fang et al., 2020), whereas the model predicted no change. The model’s value was 1.04 (Supplementary Materials, Table S7), an increase that falls just below the 5% threshold we used to define a change. Pathway length alone does not account for the small magnitude. Intermediate nodes between SL and *BRC1* also receive inputs from outside this pathway, and because the PSoup general equation averages over all incoming edges, the SL contribution is progressively attenuated at each step. A purely linear chain would transmit the signal undiluted, but these side connections diminish relative effects, leaving only a slight increase in BRC1 expression. The model therefore represents the response correctly, since *BRC1* transcript does indeed rise. Again, in the sorghum *phyB* mutant, the model predicted decreased bud release relative to the wild type, a decrease consistent with the direction reported experimentally (Figure 2A in (Kebrom *et al*., 2006), but the change fell within the no change band and was therefore recorded as a mismatch.

### Other Reasons

In the experiment with garden pea, WT treated with exogenous NPA showed no change due to buds already being so small whereas the model proposed an increase that could not be captures experimentally. WT pea treated with exogenous gibberellin after decapitation reported increased *bud release* relative to the decapitated control whereas the model predicted no change, because gibberellin acts downstream of *bud release* in the shoot branching network and therefore has no route to raise the node (Figure 5A in Cao et al., 2023). This is a matter of making a distinction around the definition of bud release verses subsequent growth, consistent with the Cao et al., (2023) study.

### Summary of the accuracy of the network for the testing data set

The testing accuracy of 75% (63 of 84) describes whether the network recovers the direction of change for perturbations never used in fitting, and at nodes other than the shoot branching output to which the network was fitted. The 21 mismatches fall into four groups. These are (i) topology not implemented, accounting for 13 mismatches, where the model tracks protein or product rather than transcript, or lacks a specific edge such as a transcription factor input, (ii) non-significant statistical difference between the perturbation and wild type in the biological experiment, accounting for 1 mismatch, where the reported experimental direction matched the prediction but did not reach significance, (iii) comparison with intermediate phenotypes, accounting for 3 mismatches, where the correct direction is recovered once multi-perturbation treatments are compared with the treatment carrying one fewer perturbation, as was also the case in training, and (iv) below cut off, accounting for 2 mismatches, where the predicted direction agrees with the experiment but the change does not cross the 5% threshold. The remaining 2 mismatches arise from other reasons discussed above. Groups (ii), (iii) and (iv) do not reflect errors in the wiring, since in each the model recovers the biological direction and the mismatch arises from the experimental statistics, the choice of baseline, or the binning threshold. Setting these aside gives a revised accuracy of 82% (69 of 84), with the remaining discrepancies confined to topology not implemented and to other reasons. The below cut off group is a limitation of the algebraic formulation, because when additional nodes feed into the path between the perturbed node and the measured node the signal is diluted at each step and the response can fall below 5% even though the direction is right.

Reproducing an observed output does not guarantee that a model’s internal intermediate states are correct, as demonstrated for parameterised network models where observables fit well while internal states remain undetermined (Raue *et al*., 2011). Although the networks used here are not parameterised in that way, the concern still applies, and comparing predicted intermediate states against measured gene expression provides a direct check that the correct phenotype is reached through the correct internal behaviour. The accuracy achieved across these testing perturbations provides some confidence that this is the case, since agreement extends to values at intermediate node states as well as to the endpoint phenotype. Reaching the correct phenotype through the correct intermediate behaviour is harder to achieve by compensating errors than reaching the endpoint alone.

Training fit is occasionally recoverable by slight rewiring, but predictive accuracy is notTo test whether the model’s accuracy depends on its curated wiring, we compared the curated network with 1000 randomised networks under each of four schemes (see Methods), simulated and scored on exactly the same perturbations, in the same way, as the curated model. The four schemes differ in how much of the curated wiring they leave intact. Sign shuffling leaves every connection exactly where it is and only swaps around which connections activate and which inhibit. Signed-degree-preserving rewiring moves the connections, but keeps for every node the number of connections in and out and the balance of activating and inhibiting ones. Degree-preserving rewiring keeps only the number of connections in and out of each node. Erdős–Rényi rewiring keeps only the nodes and the number of connections and joins them at random; it retains almost nothing of the curated network. Not one of the 4000 randomised networks matched the curated network on the training data (85%) or on the testing data (75%), under any scheme (P < 0.001; Supplementary materials, Fig. S4; Table S11). The closest any single randomised network came was 83% on the training data and 71% on the testing data, and both were sign-shuffled networks, but they were not the same network: no randomised network was among the best on both datasets at once. Averaged over its 1000 networks, sign shuffling was also the most accurate scheme on the testing data, at 48% of perturbations predicted in the correct direction (Fig. S4).

A randomised network rarely fits the training data even with an accuracy which is below the curated network. Given a training accuracy of 70% as a cut-off, sixteen percentage points below the curated network, 58 of 1000 networks met the cut-off under the scheme that changes least (sign shuffling), 24 of 1000 under signed-degree-preserving rewiring and 14 of 1000 under degree-preserving rewiring, while no Erdős–Rényi network reached 70% at all (Table S13). The networks that did fit were mostly the least randomised ones, since sign shuffling leaves every connection in place (Fig S5b).

On the occasions when a randomised network did fit the training data at 70% or above, that fit did not carry over to predict the testing dataset. The ten best-fitting networks of each scheme, selected on training accuracy alone, lost between 21 and 37 percentage points on the testing set (Table S13), whereas the curated network lost only 11 (86% to 75%). Comparing networks that fitted the training data equally well (70–80%), mean testing accuracy fell in step with how much of the curated wiring the scheme had left unchanged, at 52%, 46% and 37% against 75% for the curated network (Fig. S5c,d; Table S13), and only one of the 4000 randomised networks came within 10 percentage points of the curated network on both datasets. Taken together, these results show that the extent of randomisation determines how much predictive signal survives. Full randomisation under the Erdős–Rényi scheme performed worst, and no randomised network of this type exceeded 70% accuracy, a threshold that the more constrained randomisation schemes were able to reach by chance. Reaching it, however, did not imply predictive capability, because those same networks returned low accuracy for the testing data set which includes comparison to other nodes than *shoot branching*. The curated topology is therefore not merely fitted to the data used to build it. It retains predictive capacity in components of the network that it was never trained on. Also, it shows that each component of topology (edge, direction, sign) is important for accuracy of prediction.

## Discussion

The PSoup method used here makes several achievements that distinguish it from existing modelling frameworks for mechanistic networks; (i) it provides a transparent and reproducible method for translating accumulated biological knowledge into a testable predictive model for known perturbations in the literature, without requiring dedicated experiments for parameterization. (ii) It draws on an evidence base spanning diverse sources of information across multiple publications, laboratories, years, data types, experimental approaches, and species. (iii) It predicts change relative to a baseline condition, much the same as in developmental biology and physiology experiments, making its outputs directly comparable to biological observations. By bringing together all of this diverse data into a collective single model it ensures that the hypotheses implemented in one area of the network are sufficiently robust as to not diminish accuracy in another areas – knowledge can become holistic.

PSoup has previously been shown to recapitulate the direction of perturbation outcomes in diverse data sets for a small network model of shoot branching with 6 nodes and 7 edges (Fortuna *et al*., 2026). Here we applied the same approach to a network of the same trait built at a much larger scale, with 32 nodes and 56 edges. This shows that the method can be applied successfully to a more complex network and used to recapitulate the directional outcome of perturbations that were not used to fit the topology.

The model matched the experimental direction in 67 of 78 perturbations during training, an accuracy of 86%, subject to the caveats set out in the summary of training accuracy, and only a few of those mismatches were attributable to topology. In testing, the model matched the experimental direction in 63 of 84 perturbation-node pairs, an accuracy of 75%. Most of the testing mismatches fall outside what the model was built to represent, or follow from choices in the modelling framework such as the threshold applied to the difference between the perturbed value of a node and its baseline (see Summary of testing accuracy of the network for the remaining reasons). Setting these aside gives a revised accuracy of 81%. The headline percentage is therefore less informative than the composition of the mismatches, because those arising from framework choices place no constraint on the wiring, whereas each of the remaining mismatches identifies a specific regulatory connection absent from the current topology and could be implemented in a future version of the network. The strong performance of the model against the testing data set is indicating good generalisation of the model because, not only was the testing data set unique from the training set, it contained testing nodes from within the network that were not used for assessing outputs during training.

PSoup addresses two problems at once. Every edge is derived from a reported perturbation rather than inferred from correlation, as in most network inference methods (Maizels & Briscoe, 2026; Marbach *et al*., 2010). Every node represents a functional entity. Nodes that represent a gene, protein, metabolite, hormone pool, signalling state, phenotype or biological process can all sit in the same network at whatever level of abstraction the evidence supports. Therefore, PSoup builds networks that are causal by construction and coarse grained by design. This is close to what Maizels & Briscoe (2026) argue is needed. They describe a tension in which classical single gene knockouts provide strong causal evidence of gene function but ignore the interconnected reality of developmental systems, while network-based approaches capture that context yet increasingly supply only correlational evidence with little explanatory power. Their proposal is to treat sets of molecular components as larger functional units and abstract the molecular detail away, on the grounds that coarse graining does more than remove extraneous information, since higher levels of analysis can expose design principles and functional structure that are invisible at the level of individual interactions. They also reframe what counts as network behaviour, arguing that rather than tracking component dynamics through time, a systemic description should map the inputs of the system onto its outputs and ask what the simplest model is that reproduces those outputs. The difficulty they identify is doing this in a flexible and generic way rather than rebuilding the approach for every system.

PSoup represents an attempt to fulfill these requirements. The algebraic formulas are generated automatically from the topology rather than specified by hand, so the same procedure applies to any network assembled this way, and the model returns precisely an input to output map, since a perturbation applied at one node yields the direction of change at every other. Similarly, correlation networks do not by themselves infer causality (Krouk *et al*. 2013). PSoup encodes causality, since each edge is derived from an experiment in which a node was removed, added or blocked and the consequence recorded. Here we apply PSoup to the complex trait of shoot branching, where the network recapitulated perturbation direction with high accuracy, indicating that a coarse grained, parameter free model of this kind is sufficient to recapitulate perturbations it was never fitted to.

To test further whether the coarse-grained topology is what carries the regulatory logic, we randomised the network 1000 times (see Methods). No randomised network reproduced the model’s training or testing accuracy (Supplementary Materials, Fig. S4; Table S11). This is a standard control, since if a network of the same size and connectivity performs equally well once its edges are shuffled, the result says nothing about the biology. Networks that outperform their randomisations are instead taken to carry non-random structure (Milo *et a*l., 2002), and it is the arrangement of connections, beyond how many each node has, that gives biological networks their functional organisation and robustness (Maslov & Sneppen, 2002).

The same control has been applied to a curated plant network, where a Boolean model of the Arabidopsis cell cycle held its cyclic attractor in 88% of random perturbations against a mean of 24% across 1000 randomised networks of similar structure (Ortiz-Gutiérrez *et al*., 2015). This Boolean model and what it scored was dynamical robustness rather than directional prediction, so the test differs from ours, but the logic of the control is the same.

Further improvement of the rules based approach here exemplified by using PSoup would include iterative refinement of the network topology through systematic edge addition, removal, or sign modification, thereby identifying alternative network structures that might improve agreement with the observed biological data and therefore identify areas for future biological discovery (Krämer *et al.*, 2014).

### Implications

Molecular networks are rarely used for crop improvement (Kundu & Tanti, 2026; Leong *et al*., 2025), although several studies (Benes *et al*., 2020; Chapman, 2008; G. Hammer *et al*., 2006; Leong *et al*., 2025; Powell *et al*., 2026; Yin *et al*., 2018) have set out how they could be. Regulatory networks have been proposed as a route to guiding crop improvement (Kundu & Tanti, 2026; Leong *et al*., 2025) and integrating them into multiscale models has been argued to extend that potential further (Benes *et al*., 2020). This matters especially under a changing climate, where genotypes in breeding programmes must be assessed in environments for which no data yet exist, so that mechanistic multiscale models are needed rather than models fitted within the range of what has already been observed (Peng *et al*., 2020; Powell *et al*., 2026).

Coupling crop growth models to genomic prediction has the potential to explore genotype by environment interactions (GxE) outside of the training dataset and potentially for future climates (Cooper *et al*., 2005, 2016, 2023; Messina *et al*., 2018; Onogi, 2022; Technow *et al*., 2015).The reason of this coupling is that the crop model carries the ecophysiology, so genotype by environment interactions emerge from simulated physiology rather than being fitted as a term in statistical methods (Messina *et al*., 2018; Voss-Fels *et al*., 2019). Because training estimates biological parameters rather than environment specific effects, the crop model can then be run forward to evaluate genotypes that were never grown in the field, extending prediction from the handful of genotype by environment combinations present in the training data towards the target population of environments (Messina *et al*., 2018). That is one property of this method that makes the approach attractive for breeding under conditions that have not yet occurred.

However, the phenotype of the intermediate traits that the quantitative trait loci are connected with still sits at a considerable phenotypic distance from the cellular scale (Hammer *et al*., 2019). Placing a causal molecular network at that layer would close the distance (Hammer *et al*., 2006; Marjoram *et al*., 2014; Yin *et al*., 2018). The gap is not whether such a network belongs between markers and crop model intermediate traits, which has been argued for repeatedly, but how a network of that kind could be described in the first place and then connected with crop growth models as it is not straightforward (Hammer *et al*., 2006; Leong *et al*., 2025).

The network developed here was assembled from the available genetic and molecular physiology of shoot branching and spans several organisational levels rather than stopping at the cell, which is what allows it to connect to the organ and process scale at which the genotype specific parameters of crop models such as APSIM operate (Hammer *et al*., 2010, 2023). A causal topology changes the effect of a mechanism because the wiring itself determines how a genetic perturbation propagates to a trait (Cooper *et al*., 2002). It is hoped, that when the topology is biologically correct, the genotype-to-phenotype relationship will arise from identified mechanisms rather than fitted parameters, allowing the model to generalise to new allele combinations and environments. In this way, epistatic G×G interactions are explicitly represented to move beyond the infinitesimal model that underpins the breeder’s equation (Powell *etal.*, 2026). Getting the topology right is the prerequisite for everything downstream. Testing the network against perturbations that were not used in training, and comparing it with randomised networks to see whether they could recapitulate those same responses, provides evidence (Fig.3; Fig. 4) that the topology is a plausible representation of the prior knowledge underlying shoot branching.

That the network was assembled from species other than the crops of interest is not disqualifying. A topology grounded in conserved orthologues and conserved mechanistic networks provides a defensible scaffold onto which species specific detail can later be added. The Arabidopsis flowering network was mapped onto its maize orthologues in this way to model the floral transition of the maize shoot apex (Dong *et al*., 2012). Because the branching network was built from conserved components and performed consistently across the species on which it was tested, it offers a reasonable starting point for translation to crops such as maize or sorghum.

A further application is to supply the network as a genotype to phenotype structure for graph based genomic prediction. Tomura *et al*. (2026) tested graph attention networks for flowering time in two maize nested association mapping populations, comparing an infinitesimal structure with no marker interactions, a fully connected structure, and a data-driven prior knowledge structure in which edges were inferred from the data by a random forest with Shapley scores. The prior knowledge model did not consistently outperform the others, although an ensemble of all three did. That prior knowledge was data-driven and therefore bounded, since it was inferred from the same dataset it was then used to predict and carries only the interactions expressed in those populations and environments, so the result is not a test of the value of biological prior knowledge derived independently. Because a PSoup network is constructed from published perturbation experiments rather than inferred from marker–phenotype associations, it is not bounded by the population or environments used for training. Supplying it as the graph structure for a graph-based model therefore tests whether causal structure established independently of the breeding dataset improves prediction.

The predictions from a PSoup network can be used to search expression data for genes of an as yet undiscovered function. For example, a network can predict complex co-expression patterns derived from a number of varied experimental constructs. The LBO mutant was discovered in this way by screening transcriptome data from eight differing experimental constructs (wild type, strigolactone mutants, decapitation and auxin treatment) (Brewer *et al*., 2016). In this example the *LBO* gene showed a complex gene expression pattern matching that of *MAX3*, that was then confirmed by reverse genetics. We can imagine using this principle at scale for more complex and widespread discovery by analysing diverse expression patterns in transcriptomic data without relying on established patterns of gene expression.

Beyond agriculture, these networks can be used to improve experimental design and data interpretation. One can imagine a near future where AI assists model development and its iteration through experimental cycles in lab and field, continually improving knowledge. For example, this brings us to a point where we can start to use complex network topology to interrogate transcriptomic and other highly comprehensive data sets in a manner moving far along from correlative and associative approaches.

## Limitations

The model predicts whether a perturbation will increase or decrease branching relative to wild type but cannot quantify the magnitude of change. If magnitude matters, node weights must be calibrated against quantitative biological data. The model implementation can be subjective around how to implement the wide range of possibilities of biological interactions and associated perturbations. Because the model carries no edge strengths, the influence of a perturbation is shaped primarily by its position in the network. An effect originating far from the output trait is attenuated as it propagates through intermediate nodes that are themselves regulated by others. In biology, however, a distant gene can exert a strong effect on the trait, which the model could capture only if weights were assigned to amplify that node’s contribution. Additionally, the model relies heavily on prior knowledge; when that knowledge is weak or incomplete, network accuracy is compromised. However, once a robust topology is attained using an approach such as outlined herein, statistical approaches aimed at fitting parameters to phenotypic data could resolve this.

The model remains limited in areas where biological understanding is incomplete or conflicting, particularly in SMXL, ABA and auxin-related regulatory mechanisms, highlighting the need for additional experimental data to refine these network regions. The model is predictive only for perturbations reported in the literature, and it remains unknown whether it can predict novel perturbations that are not described there. It should therefore be used with caution, and its outputs should be treated as hypotheses to be tested rather than as established predictions.

Finally, the manual curation required here was painstaking and slow. Reaching the final model took 89 iterations with every step handled by hand, which highlights the difficulty of building a regulatory network for even a single trait. This is precisely the kind of process that agentic AI could assist, enabling faster network construction and scaling the approach well beyond what manual curation allows (Mitsanis *et al*., 2026).

## Conclusion and Future Directions

This study shows that a directed, signed network built from prior biological knowledge of shoot branching and grounded in diverse scientific data, once translated into algebraic formulas by PSoup, can represent effects of genetic and other perturbations and on phenotypes including plant architectural (branching and bud outgrowth) and molecular traits (gene expression). The model recapitulates the direction of change in 75% of 84 perturbation testing node pairs withheld from training (which was 86% accurate). Out of 4000 randomised networks none compared with this accuracy. Therefore, prior knowledge of a system as expressed in a network topology is alone able express the regulatory logic underpinning the trait, offering a way to synthesise highly diverse experimental results into a model that predicts knowledge it was never fitted to. It also shows that PSoup, previously tested only on a small network (Fortuna *et al*., 2026), can be used for a more complex network. This paves the way for innovation in crop breeding as well as in approaches to better understand plants as systems. The main constraint is manual curation, which was slow and painstaking and limits how quickly the approach can be extended to other traits. Automating this step with agentic AI is our next step (Mitsanis *et al*., 2026).

## Acknowledgments

We would like to thank the members of the Australian Research Council (ARC) Centre of Excellence for Plant Success in Nature and Agriculture (CE200100015), many of whom contributed valuable feedback during the creation and testing of the network. We also thank David Kainer for the design style of Figure 2 (created in Cytoscape);. The project was also supported by an ARC Georgina Sweet Laureate Fellowship (FL180100139).

## Data Availability

The code and data for this study are available at https://github.com/CMits/Shoot_Brancing_PSoup.git.

## Notes

https://github.com/CMits/Shoot_Brancing_PSoup.git

### Competing Interest Statement

The authors have declared no competing interest.

