## Supplementary Information for "From Diverse Prior Knowledge to Mechanistic Causal Network Using PSoup: A Case Study in Shoot Branching"

### Supplementary Materials

#### Supplementary Figures

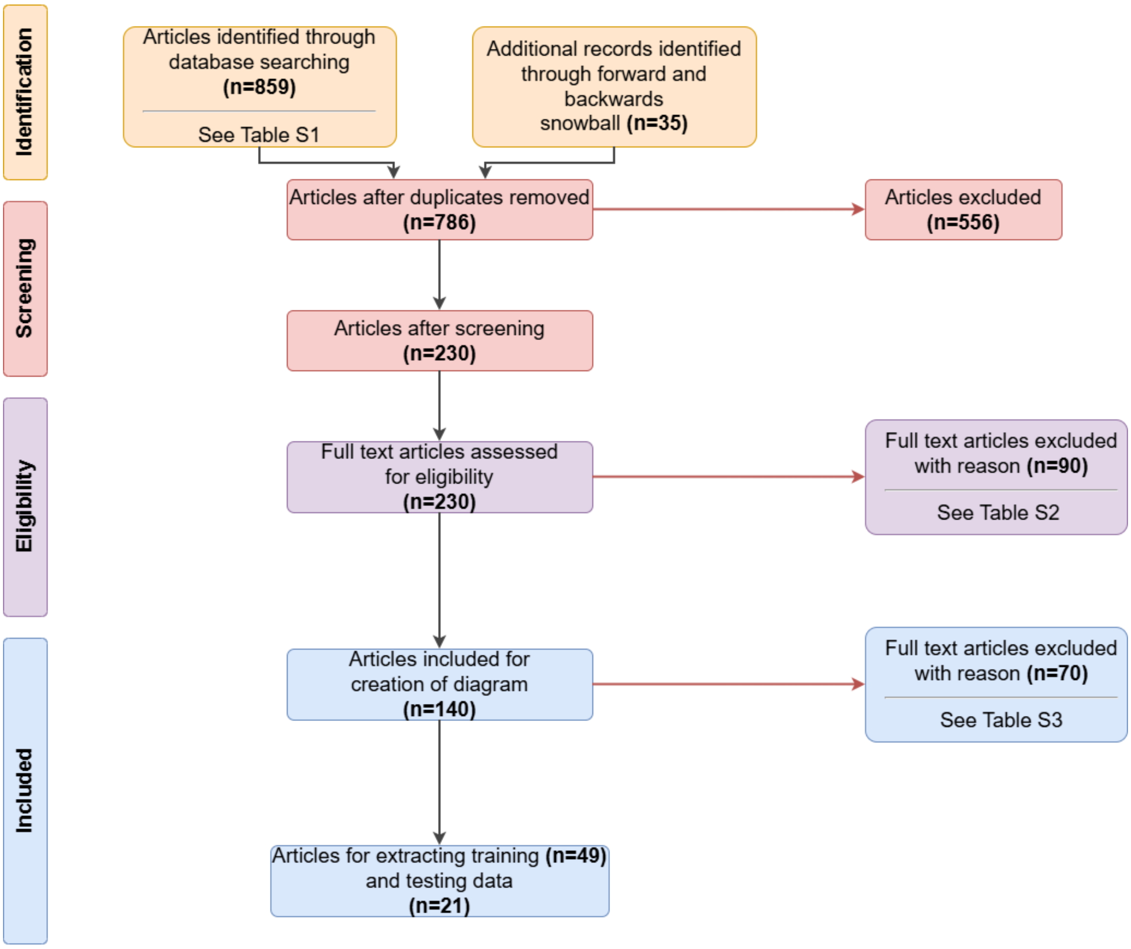


Fig. S1. PRISMA flow diagram illustrating the systematic literature review process used to identify relevant studies for constructing the shoot branching signaling network and extracting training and testing datasets. Three Scopus searches (Table S1) combined with forward and backward snowballing yielded 894 articles, of which 786 remained after duplicate removal. Articles were screened for eligibility using the criteria listed in Table S2 (n=140 remained; Dataset S1) and subsequently filtered for data extraction suitability using the criteria in Table S3 (n=70 remained, from which 49 provided quantitative data for model training and 21 for testing; Datasets S2 and S3). The domain analysis was performed from the years 2002-2024.

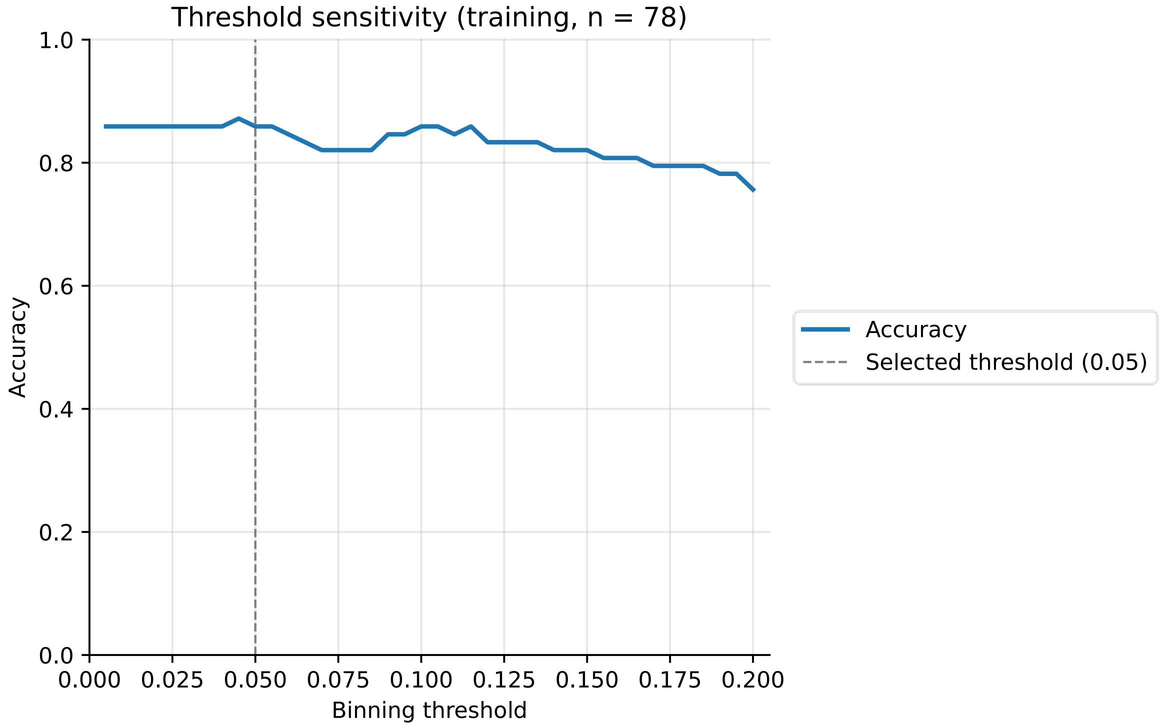


Fig. S2. Threshold sensitivity analysis. For each candidate binning threshold between 0.005 and 0.200 the ratio of perturbation to control Shoot Branching values was discretised into three bins (increase, decrease or unchanged) and the resulting predictions were compared with the literature-curated biological bins across the 78 training comparisons in Dataset S9. The solid line shows the resulting accuracy and the vertical dashed line marks the threshold of 0.05 used throughout the rest of the analysis. Accuracy sits on a plateau at low thresholds and declines progressively beyond 0.05 as stricter cut-offs force more comparisons into the increase or decrease categories.

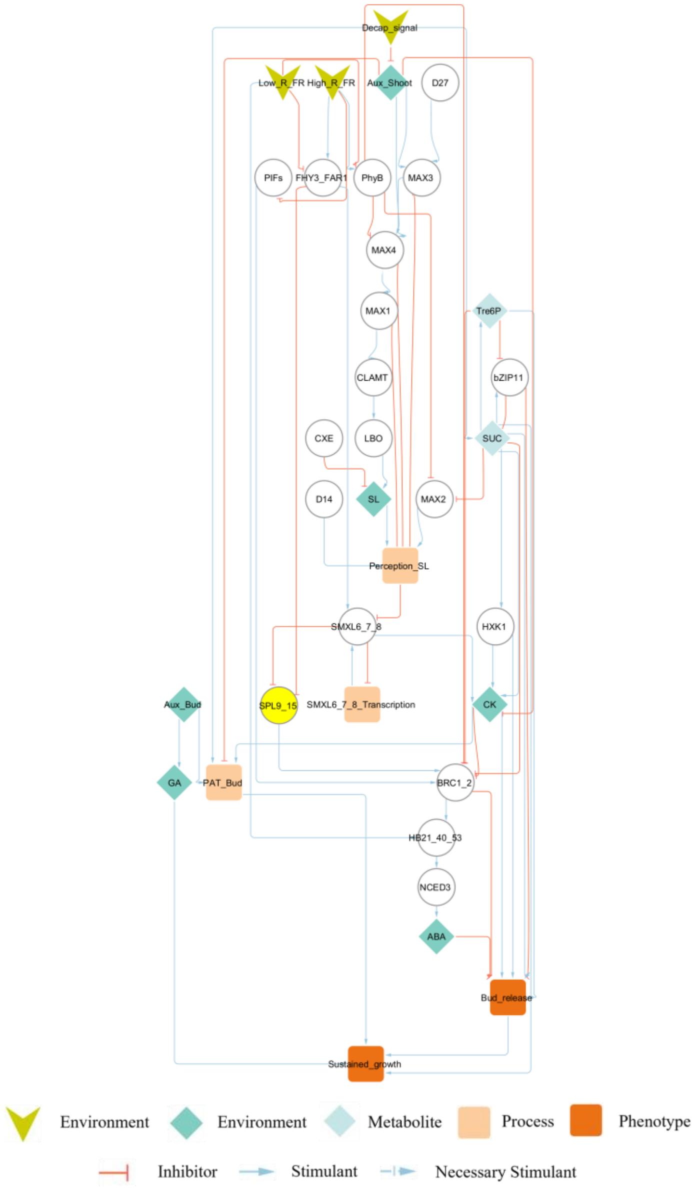


Fig. S3. Expanded shoot branching network integrating sucrose signalling. The network contains 35 nodes (the 32 nodes of the original model plus three new sucrose pathway nodes Tre6P, HXK1 and bZIP11) and 66 directed edges (40 stimulatory and 26 inhibitory). Green arrows indicate stimulatory interactions and red arrows indicate inhibitory interactions. Ten new edges connect the sucrose signalling nodes to the existing network, so that SUC stimulates Tre6P, HXK1 and bZIP11, Tre6P inhibits BRC1/2 and bZIP11 and stimulates Bud_release, HXK1 stimulates CK and Bud_release, and bZIP11 inhibits Bud_release and SUC. The new edges come from Fichtner *et al*. (2021) for Tre6P, Barbier *et al*. (2021) for HXK1 and Kreisz *et al*. (2024) for bZIP11. The expanded network reaches an accuracy of 64 of 78 (82.05%) on the 78 training perturbations reported in Dataset S14, 11 of 11 (100%) on the eleven additional sucrose perturbations listed in Table S9, and 62 of 84 (73.81%) on the 84 testing perturbations reported in Dataset S15. The network is depicted in Cytoscape using a hierarchical layout.

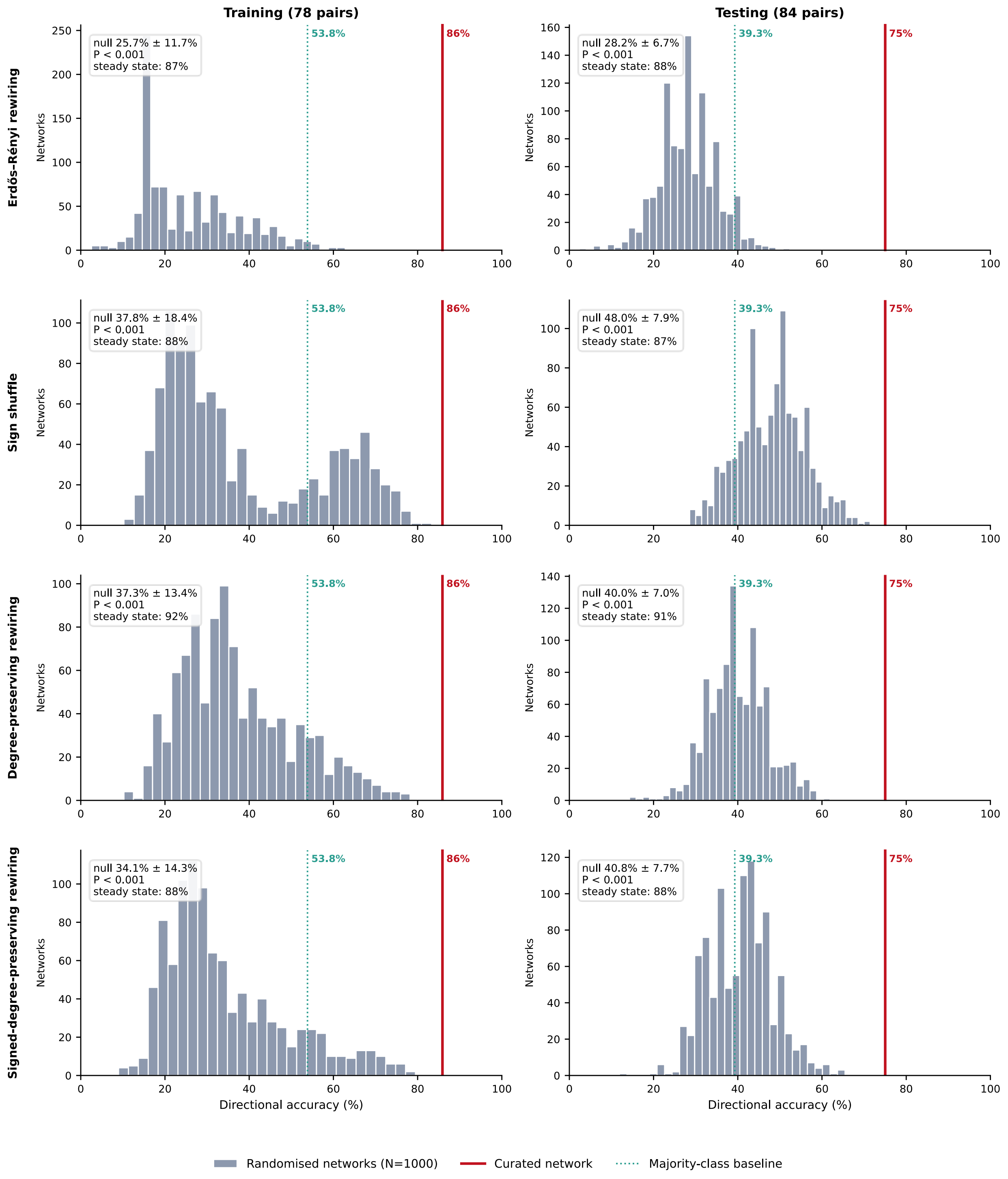


Fig. S4. Comparison of the curated shoot branching network against randomised networks directional accuracy across perturbations. Each panel shows the distribution of directional accuracy across 1000 randomised networks (grey), the accuracy of the curated topology (red line) and the majority-class baseline (green dotted line), for the training (left panels) and testing (right panels) sets. The four network-randomisation families are ordered by how much structure they hold fixed, from sign shuffling, which leaves every edge in place and changes only their signs, to signed-degree-preserving rewiring, which preserves every node's in-degree, out-degree and balance of activating and inhibiting connections and destroys only the specific wiring. Networks that failed to reach a steady state were scored as incorrect rather than excluded. Inset values give the mean and standard deviation of the randomised distribution, the empirical P value, and the proportion of randomised networks reaching a steady state. The randomisations differ substantially from the curated network: sign shuffling leaves every edge in place and changes a mean of 21.9 of the 52 signs, whereas the three rewiring schemes displace a mean of 42.9 to 49.6 edges and leave only 5.3 (degree-preserving), 8.2 (signed-degree-preserving) or 0.9 (Erdős–Rényi) of the 52 curated edges in place. Networks that differ from the curated one by only a few edges are examined separately in Fig. S5 and Table S12. The green dotted line in each panel marks the majority-class baseline: the accuracy obtained by ignoring the network entirely and assigning every perturbation to the single most frequent observed outcome. In the training set that outcome is an increase in branching, observed for 42 of the 78 perturbations (53.8%); in the testing set decreases and increases are equally common, at 33 of the 84 perturbations each (39.3%). The baseline is the floor that any informative model must clear, and it is the reference against which the randomised distributions should be read: an ensemble sitting at or below the line is behaving as an uninformative classifier, which is the expected result, whereas an ensemble sitting well above it would mean the observed directions are partly predictable from class frequencies alone and the topology claim would need qualifying. The curated network clears the baseline by about 32 percentage points on the training set (67 of 78, 86%, against 53.8%) and by about 36 points on the testing set (63 of 84, 75%, against 39.3%). Because the two datasets differ in class balance the baseline differs between the left and right columns, so each panel should be read against its own baseline and its own randomised distribution rather than across columns — the same point made for Fig. S5b.

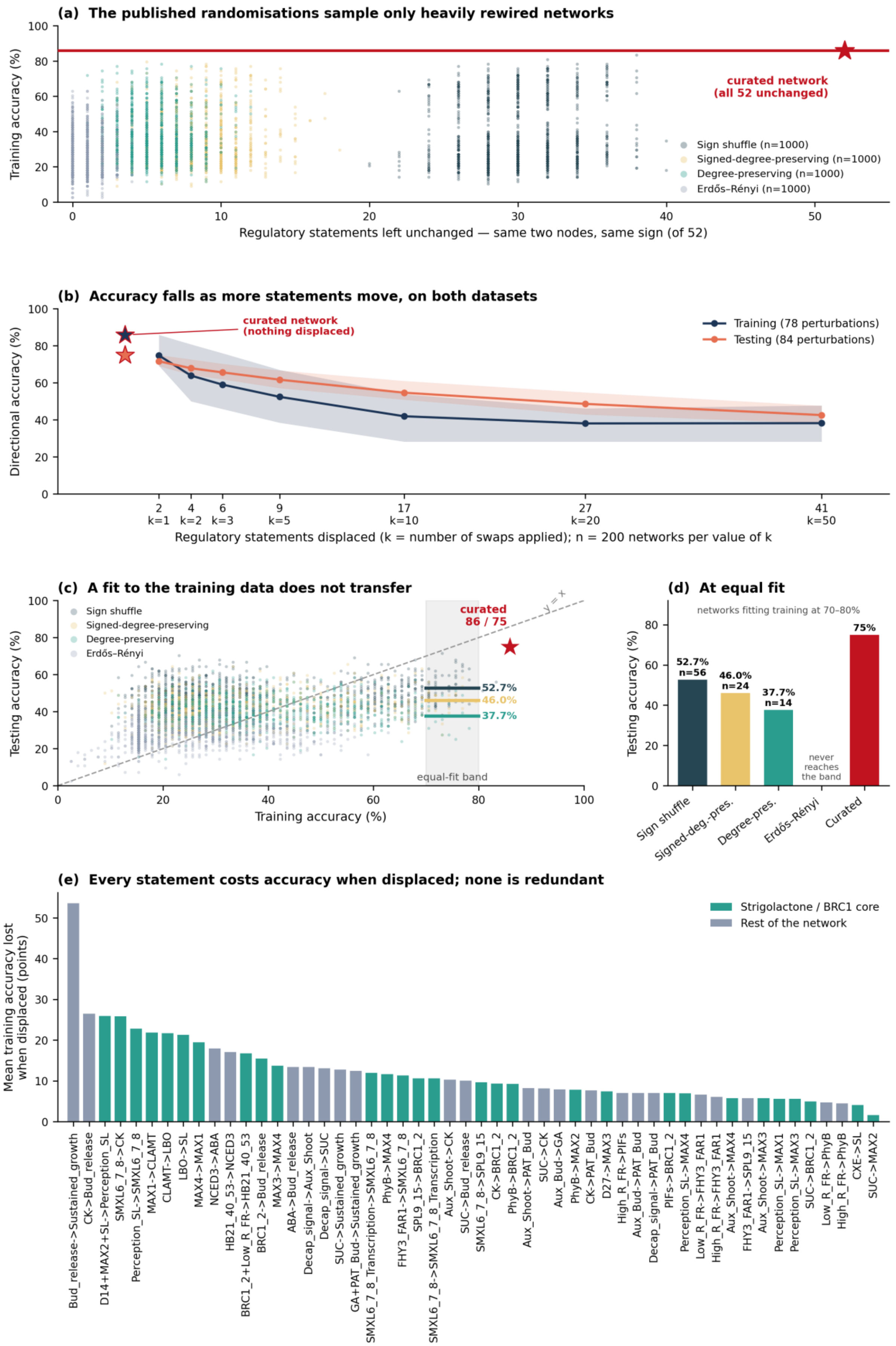
(a) The published randomisations sample only heavily rewired networks
Training accuracy (%)
curated network
(all 52 unchanged)
Sign shuffle (n=1000)
Signed-degree-preserving (n=1000)
Degree-preserving (n=1000)
Erdős–Rényi (n=1000)
Regulatory statements left unchanged — same two nodes, same sign (of 52)
(b) Accuracy falls as more statements move, on both datasets
Directional accuracy (%)
curated network
(nothing displaced)
Training (78 perturbations)
Testing (84 perturbations)
2 k=1
4 k=2
6 k=3
9 k=5
17 k=10
27 k=20
41 k=50
Regulatory statements displaced (k = number of swaps applied); n = 200 networks per value of k
(c) A fit to the training data does not transfer
Testing accuracy (%)
Sign shuffle
Signed-degree-preserving
Degree-preserving
Erdős–Rényi
curated
86 / 75
y = x
52.7%
46.0%
37.7%
equal-fit band
Training accuracy (%)
(d) At equal fit
networks fitting training at 70–80%
Testing accuracy (%)
52.7%
n=56
46.0%
n=24
37.7%
n=14
never
reaches
the band
75%
Sign shuffle
Signed-deg.-pres.
Degree-pres.
Erdős–Rényi
Curated
(e) Every statement costs accuracy when displaced; none is redundant
Mean training accuracy lost
when displaced (points)
Strigolactone / BRC1 core
Rest of the network
Bud_release->Sustained_growth
CK->Bud_release
D14+MAX2+SL->Perception_SL
SMXL6_7_8->CK
Perception_SL->SMXL6_7_8
MAX1->CLAMT
CLAMT->LBO
LBO->SL
MAX4->MAX1
NCED3->ABA
HB21_40_53->NCED3
BRC1_2+Low_R_FR->HB21_40_53
BRC1_2->Bud_release
MAX3->MAX4
ABA->Bud_release
Decap_signal->Aux_Shoot
Decap_signal->SUC
SUC->Sustained_growth
GA+PAT_Bud->Sustained_growth
SMXL6_7_8_Transcription->SMXL6_7_8
PhyB->MAX4
FHY3_FAR1->SMXL6_7_8
SPL9_15->BRC1_2
SMXL6_7_8->SMXL6_7_8_Transcription
Aux_Shoot->CK
SUC->Bud_release
SMXL6_7_8->SPL9_15
CK->BRC1_2
PhyB->BRC1_2
Aux_Shoot->PAT_Bud
SUC->CK
Aux_Bud->GA
PhyB->MAX2
CK->PAT_Bud
D27->MAX3
High_R_FR->PIFs
Aux_Bud->PAT_Bud
Decap_signal->PAT_Bud
PIFs->BRC1_2
Perception_SL->MAX4
Low_R_FR->FHY3_FAR1
High_R_FR->FHY3_FAR1
Aux_Shoot->MAX4
FHY3_FAR1->SPL9_15
Aux_Shoot->MAX3
Perception_SL->MAX1
Perception_SL->MAX3
SUC->BRC1_2
Low_R_FR->PhyB
High_R_FR->PhyB
CXE->SL
SUC->MAX2

Fig. S5. Networks that differ from the curated network by only a few regulatory statements, and whether a fit to the training data transfers. Throughout, a statement counts as unchanged only if it runs between the same two nodes and carries the same sign; Table S13 breaks this down. (a) Each point is one of the 4000 randomised networks summarised in Fig. S4, plotted by how many of the 52 statements it leaves unchanged against its training accuracy; the curated network is the star at 52. The schemes leave a mean of only 0.9 to 8.2 statements unchanged, so they populate only the heavily rewired part of the axis and do not test networks close to the curated one. (b) Accuracy against the number of statements displaced, using the degree-preserving swap with a budget of k = 1 to 50 swaps and 200 networks per value of k, for the training set (navy) and the testing set (orange). Lines show the mean and shaded areas the interquartile range; stars mark the curated network, which has nothing displaced. Accuracy falls as soon as a few statements move, on both datasets. The testing curve lies above the training curve once several statements have moved, because the testing set has a different class balance and so returns a higher accuracy for an uninformative network; each dataset should therefore be compared against its own randomised distribution (Table S11) rather than against the other. (c) Testing against training accuracy for the 4000 randomised networks, with a y = x reference. Points lie well below the diagonal, so a given level of fit to the training data does not reappear on the held-out data. The shaded band marks networks that fit the training data equally well (70% to 80% training accuracy); the horizontal line within it gives each scheme's mean testing accuracy in that band. (d) The same equal-fit comparison as bars, with the number of networks contributing to each. Testing accuracy falls from 52.7% for sign shuffling, which leaves every connection in place, to 46.0% and 37.7% for the two rewiring schemes, in step with how much wiring each destroys; no Erdős–Rényi network reaches the band at all, and the curated network reaches 75%. (e) The mean training accuracy lost when each of the 52 statements is displaced, from the exhaustive enumeration of all 1016 admissible single swaps. Every statement costs accuracy when displaced, so none is redundant with respect to the data. Bars are coloured by whether the statement touches the strigolactone or BRC1 core. Source data: Datasets S9 and S13; per-statement values in Table S12; scheme summaries in Table S13.

### Supplementary Tables

Table S1. Scopus search strings used for the systematic literature review. The domain analysis was performed in February 2024, and it covers literature from 2002.

| **Query string** |
| --- |
| "shoot branching" AND (mutant OR knockout OR overexpression) AND (Arabidopsis OR rice OR pea) |
| "tiller number" AND (mutant OR treatment) AND (sorghum OR rice OR wheat) |
| "bud outgrowth" AND (strigolactone OR cytokinin OR auxin) |

Table S2. Paper exclusion criteria used during the screening process in the PRISMA figure S1 for papers to extract information for the Shoot Branching Knowledge graph. The domain analysis was performed in February 2024, and it covers literature from 2002.

| **Exclusion criterion** |
| --- |
| Articles without full text available |
| Articles not written in English |
| Duplicate publication |
| Publications that are not articles (e.g., survey) |
| Articles do not associate with shoot branching |
| Articles do not associate with hormones and shoot branching |
| Articles not relevant with the focus of the study |
| Articles published before 2002 |

Table S3. Paper exclusion criteria used during the screening process for the training and testing dataset. The domain analysis was performed in February 2024, and it covers literature from 2002.

| Exclusion criterion |
| --- |
| Paper does not report shoot branching, tillering, or bud outgrowth phenotypes |
| Study conducted in woody species (trees) |
| No mutant or treatment comparison to wild type |
| Quantitative data insufficient for qualitative interpretation |
| Review paper without original experimental data |

Table S4. Treatments excluded from the training and testing sets for lack of biological consensus. Of the 84 unique training perturbations (Dataset S4) and 88 unique testing perturbation–node pairs (Dataset S5), 4 training treatments and 4 testing pairs were excluded, giving consensus sets of 80 training (78 retained after excluding two erroneous export records) and 84 testing comparisons, because their reported biological outcomes conflicted across independent experiments or rested on a single experiment. Columns, left to right: Set (training or testing); Condition (the perturbation, i.e. treatment or genotype combination); Compared to (the baseline used); Node measured (the network node scored); Species; Supporting publications (each source with the direction it reported and the species tested); and Reason for exclusion. Directions are coded as increase (+1), unchanged (0) or decrease (−1) relative to control. Mutants are written as perturbations to nodes in the form GENE_n.

| Set | Condition | Compared to | Node measured | Species | Supporting publications | Reason for exclusion |
|---|---|---|---|---|---|---|
| Training | BRC1_2_n, +SL | BRC1_2_n | Shoot Branching | Arabidopsis, Pea, Rice | (Seale *et al.*, 2017) — Decrease (Arabidopsis); (van Rongen *et al.*, 2019) — Decrease (Arabidopsis); (Brewer *et al.*, 2009) — Unchanged (Pea); (Minakuchi *et al.*, 2010) — Unchanged (Rice) | 50:50 split between Decrease and Unchanged across 4 experiments — no majority reaches the consensus threshold. |
| Training | CXE_n | WT | Shoot Branching | Arabidopsis | (Roesler *et al.*, 2021) — Increase (Arabidopsis); (Xu *et al.*, 2021) — Increase (Arabidopsis); (Xu *et al.*, 2021) — Unchanged (Arabidopsis) | 33% Unchanged vs 66% Increase across 3 experiments — no bin reaches the 50%+ consensus majority. |
| Training | HB21_40_53_n | WT | Shoot Branching | Arabidopsis | (González-Grandío *et al.*, 2017) — Unchanged (Arabidopsis); (González-Grandío *et al.*, 2017) — Increase (Arabidopsis) | 50:50 split between Unchanged and Increase across 2 experiments — no majority reaches the consensus threshold. |
| Training | PIN_1_3_4_7_n, MAX2_n | WT | Shoot Branching | Arabidopsis | (Bennett *et al.*, 2016) — Unchanged (Arabidopsis); (van Rongen *et al.*, 2019) — Increase (Arabidopsis) | 50:50 split between Unchanged and Increase across 2 experiments — no majority reaches the consensus threshold. |
| Training | SMXL6_7_8_n | WT | Shoot Branching | Arabidopsis, Rice | (Fichtner *et al.*, 2022) — Unchanged (Arabidopsis); (Seale *et al.*, 2017) — Increase (Arabidopsis); (Song *et al.*, 2017) | Three-way split (2/1/4 for Decrease/Unchanged/Increase) across 7 experiments across 2 species — reflects species-specific or context-dependent biology, no clear majority. |

| | | | | | | |
|---|---|---|---|---|---|---|
| | | | | | — Increase (Rice); (Wang *et al.*, 2015) — Decrease (Arabidopsis); (Xie *et al.*, 2020) — Decrease (Arabidopsis); (Zhou *et al.*, 2013) — Increase (Rice); (Fang *et al.*, 2020) — Increase (Rice) | |
| Testing | Decap_signal_n, +SL | Decap_signal_n | Bud_release | Pea | (Cao *et al.*, 2023) — Decrease (Pea); (Brewer *et al.*, 2009) — Unchanged (Pea) | 50:50 split between Decrease and Unchanged across 2 experiments — no majority reaches the consensus threshold. |
| Testing | MAX2_n, +SL | WT | Bud_release | Pea | (Cao *et al.*, 2023) — Decrease (Pea); (Dun *et al.*, 2012) — Unchanged (Pea) | 50:50 split between Decrease and Unchanged across 2 experiments — no majority reaches the consensus threshold. |
| Testing | WT, +CK, +SL | WT | Bud_release | Pea | (Cao *et al.*, 2023) — Increase (Pea); (Dun *et al.*, 2012) — Decrease (Pea) | 50:50 split between Decrease and Increase across 2 experiments — no majority reaches the consensus threshold. |
| Testing | WT, +SUC, +SL | WT | Bud_release | Pea, Rice, Rosa hybrida | (Bertheloot *et al.*, 2020) — Decrease (Rosa hybrida); (Cao *et al.*, 2023) — Decrease (Pea); (Patil *et al.*, 2022) — Increase (Rice) | 66% Decrease vs 33% Increase across 3 experiments — no bin reaches the 50%+ consensus majority. |

Table S5. Training confusion matrix and per-class metrics (threshold = 0.05, n = 78). The upper block cross-tabulates the PSoup-predicted branching direction (columns) against the expected biological direction (rows) for the 78 training comparisons, with bins coded increase (+1), unchanged (0) or decrease (−1). Predicted direction was taken from the ratio of perturbation to control *Shoot Branching* values (the Sustained_growth node in the simulation datasets): increase or decrease was scored correct when the ratio rose above or fell below one, and unchanged was scored correct when the ratio stayed within 5% of one (0.95–1.05). The lower block reports per-class precision, recall, F1 and support for each direction bin. Source data: Dataset S9.

| Expected direction | −1 (Predicted) | 0 (Predicted) | +1 (Predicted) |
|---|---|---|---|
| −1 | 22 | 2 | 0 |
| 0 | 7 | 5 | 0 |
| +1 | 2 | 0 | 40 |

| Class | Precision | Recall | F1 | Support |
|---|---|---|---|---|
| −1 | 0.71 | 0.92 | 0.80 | 24 |
| 0 | 0.71 | 0.42 | 0.53 | 12 |
| +1 | 1.00 | 0.95 | 0.98 | 42 |

Table S6. Training discrepancies (5% no-change band; n = 11). The 11 training comparisons where the PSoup-predicted branching direction did not match the expected biological direction, scored as in Table S5. Columns, left to right: Perturbation (the treatment or genotype combination); Control (its baseline); Control branching and Pred. branching (the simulated Shoot Branching values for the control and the perturbation, i.e. the Sustained_growth node in the datasets); Pred. bin and Bio. bin (the predicted and biological directions of change, coded increase (+1), unchanged (0) or decrease (−1)); Species; and Supporting Publication(s). Mutants are represented as perturbations to nodes written as GENE_n. The Shoot Branching node is referred to as Sustained_growth in the underlying datasets. Pred. branching and Control branching are taken from Dataset S9. Change (%) is the predicted value relative to its control, (predicted / control - 1) x 100; a comparison is binned as no change when this lies within +/-5%, so the threshold is relative to the control rather than an absolute difference.

| **Phenotypic comparison** | | **Model Output** | | | **Bins** | | | |
|---|---|---|---|---|---|---|---|---|
| **Perturbation** | **Control** | **Control branching** | **Pred. branching** | **Change (%)** | **Pred. bin** | **Bio. bin** | **Species** | **Supporting Publication(s)** |
| BRC1_2_n, +Low_R_FR | BRC1_2_n | 2.00 | 1.91 | -4.3 | 0 | -1 | Arabidopsis | (González-Grandío *et al.*, 2017) |
| BRC1_2_n, PIN_1_3_4_7_n, +SL | BRC1_2_n, PIN_1_3_4_7_n | 1.25 | 1.19 | -5.0 | 0 | -1 | Arabidopsis | (van Rongen *et al.*, 2019) |
| FHY3_FAR1_n, +Low_R_FR | FHY3_FAR1_n | 0.81 | 0.73 | -9.7 | -1 | 0 | Arabidopsis | (Xie *et al.*, 2020) |
| HB21_40_53_n, +ABA | HB21_40_53_n | 1.33 | 1.00 | -25.0 | -1 | 0 | Arabidopsis | (González-Grandío *et al.*, 2017) |
| HB21_40_53_n, +Low_R_FR | HB21_40_53_n | 1.33 | 1.22 | -8.7 | -1 | 0 | Arabidopsis | (González-Grandío *et al.*, 2017) |
| PIN_1_3_4_7_n | WT | 1.00 | 0.62 | -37.5 | -1 | 0 | Arabidopsis | (Bennett *et al.*, 2016) |
| PIN_1_3_4_7_n, MAX4_n | WT | 1.00 | 0.81 | -19.0 | -1 | 1 | Arabidopsis | (van Rongen *et al.*, 2019) |
| SMXL6_7_8_n, +SL | SMXL6_7_8_n | 1.27 | 1.16 | -8.7 | -1 | 0 | Rice | (Song *et al.*, 2017) |
| SMXL6_7_8_n, MAX2_n | WT | 1.00 | 0.64 | -35.6 | -1 | 0 | Arabidopsis | (Seale *et al.*, 2017) |
| SMXL6_7_8_n, SPL9_15 | WT | 1.00 | 0.89 | -11.0 | -1 | 0 | Arabidopsis | (Xie *et al.*, 2020) |
| WT, +NPA | WT | 1.00 | 0.66 | -34.0 | -1 | 1 | Rice | (Lin *et al.*, 2009) |

Table S7. Testing discrepancies (5% no-change band; n = 21). The 21 unique perturbation–node combinations where the PSoup-predicted direction did not match the expected biological direction. Columns, left to right: Perturbation (the treatment or genotype combination); Compare with (its baseline); Node (the network node measured); Pred. value and Control value (the simulated node values for the perturbation and the control); Pred. bin and Bio. bin (the predicted and biological directions of change, coded increase (+1), unchanged (0) or decrease (−1)); Species; and Supporting Publication(s). Source data: Dataset S13.Mutants are represented as perturbations to nodes written as GENE_n. Change (%) is the predicted value relative to its control, (predicted / control - 1) x 100; a comparison is binned as no change when this lies within +/-5%, so the threshold is relative to the control rather than an absolute difference.

| **Phenotypic Comparison** | | | **Model Output** | | | **Bins** | | | |
|---|---|---|---|---|---|---|---|---|---|
| **Perturbation** | **Compare with** | **Node** | **Pred. value** | **Control value** | **Change (%)** | **Pred. bin** | **Bio. bin** | **Species** | **Supporting Publication (s)** |
| BRC1_2_n, +Low_R_FR | WT | NCED3 | 0.00 | 1.00 | -100.0 | -1 | 0 | Arabidopsis | (González-Grandío *et al*., 2017) |
| D14_n, Decap_signal_n | WT | MAX4 | 1.19 | 1.00 | +18.5 | 1 | -1 | Pea | (Foo *et al*., 2005) |
| D27_n | WT | MAX4 | 0.00 | 1.00 | -100.0 | -1 | 1 | Rice | (Jiang *et al*., 2013); (Zhou *et al*., 2013) |
| Decap_signal_n, +GA | Decap_signal_n | Bud_release | 2.58 | 2.58 | +0.0 | 0 | 1 | Pea | (Cao *et al*., 2023) |
| MAX2_n, Decap_signal_n | WT | MAX4 | 1.19 | 1.00 | +18.5 | 1 | -1 | Pea | (Foo *et al*., 2005) |
| MAX2_n, Decap_signal_n, +Aux_Shoot | MAX2_n, Decap_signal_n | MAX4 | 10.67 | 1.19 | +800.0 | 1 | -1 | Pea | (Foo *et al*., 2005) |
| MAX3_n | WT | MAX4 | 0.00 | 1.00 | -100.0 | -1 | 1 | Pea, Rice | (Foo *et al*., 2005); (Minakuchi *et al*., 2010) |
| MAX4_n, +CK, +SL | MAX4_n | BRC1_2 | 0.86 | 1.00 | -14.3 | -1 | 0 | Pea | (Dun *et al*., 2012) |
| MAX4_n, +SL | MAX4_n | BRC1_2 | 1.00 | 1.00 | +0.0 | 0 | 1 | Pea, Rice | (Dun *et al*., 2012); (Kerr *et al*., 2020); (Fang *et al*., 2020) |
| PhyB_n | WT | Bud_release | 0.95 | 1.00 | -4.8 | 0 | -1 | Sorghum | (Kebrom *et al*., 2006) |
| SMXL6_7_8_n | WT | MAX4 | 1.00 | 1.00 | +0.0 | 0 | 1 | Rice | (Zhou *et al*., 2013) |
| SPL9_15_n | WT | SMXL6_7_8_Transcription | 1.00 | 1.00 | +0.0 | 0 | -1 | Rice | (Song *et al*., 2017) |
| WT, +NPA | WT | Bud_release | 0.89 | 1.00 | -10.8 | -1 | 0 | Pea | (Brewer *et al*., 2009) |
| WT, +NPA | WT | D14 | 1.00 | 1.00 | +0.0 | 0 | -1 | Arabidopsis | (Chevalier *et al*., 2014) |

| | | | | | | | | | |
|---|---|---|---|---|---|---|---|---|---|
| Decap_signal_n, +NPA | Decap_signal_n | Bud_release | 2.72 | 2.23 | +22.4 | 1 | -1 | Pea | (Brewer *et al.*, 2009) |
| WT, +SL | WT | BRC1_2 | 1.04 | 1.00 | +4.4 | 0 | 1 | Arabidopsis, Pea | (Chevalier *et al.*, 2014); (Dun *et al.*, 2012) |
| WT, +SL | WT | MAX2 | 1.00 | 1.00 | +0.0 | 0 | -1 | Arabidopsis | (Chevalier *et al.*, 2014) |
| WT, +SL | WT | MAX4 | 0.69 | 1.00 | -30.8 | -1 | 0 | Arabidopsis | (Chevalier *et al.*, 2014) |
| WT, +SUC | WT | D14 | 1.00 | 1.00 | +0.0 | 0 | -1 | Rice | (Patil *et al.*, 2022) |
| WT, +SUC | WT | SMXL6_7_8_Transcription | 0.96 | 1.00 | -4.2 | 0 | -1 | Rice | (Patil *et al.*, 2022) |
| WT, +SUC, +Aux_Shoot | WT | Bud_release | 2.07 | 1.00 | +106.7 | 1 | -1 | Rosa hybrida | (Bertheloot *et al.*, 2020) |

### Supplementary Text 1

The model predicts the qualitative direction of change in shoot branching (or gene expression in the testing dataset) relative to a defined baseline. The baseline used for each comparison depends on the nature of the perturbation, following the same logic used in the original experimental studies. Three categories of baseline comparisons were used.

First, single and multiple genetic mutants without exogenous treatments were compared to wild type (WT). This is the standard comparison used in genetic studies, where the effect of a mutation is assessed relative to the unperturbed plant. In the training dataset, 40 of 78 comparisons used this baseline, and in the testing dataset, 31 of 84 comparisons did so (Dataset S9 for training, Dataset S13 for testing). For example, the max4 single mutant and the max3/max4 double mutant were both compared to WT to determine whether branching increased, decreased, or remained unchanged.

Second, when an exogenous hormone or signal was applied to a mutant background, the comparison baseline was the mutant without the exogenous treatment. This mirrors the experimental design in the source publications, where the effect of an exogenous application is measured against the untreated mutant rather than against WT, isolating the effect of the exogenous signal from the effect of the mutation. In the training dataset, 30 comparisons used this baseline, and in the testing dataset, 12 comparisons did so. For example, the max4 mutant treated with exogenous SL (max4 + SL) was compared to max4 alone, and the d14 mutant treated with exogenous auxin (d14 + auxin) was compared to d14 alone.

Third, exogenous treatments applied to WT plants were compared to untreated WT. In the training dataset, 9 comparisons used this baseline, and in the testing dataset, 29 comparisons did so. This includes both single exogenous applications (e.g., WT + SL compared to WT) and combined exogenous applications (e.g., WT + CK + SL compared to WT). For combined exogenous treatments on WT, the comparison was made to untreated WT rather than to the single exogenous treatment, consistent with how these experiments were reported in the original publications.

It is important to note that for multiple genetic mutants (e.g., max3/max4 double knockout), the standard comparison was against WT rather than against a single mutant. While this is the conventional comparison used in the original experimental studies, it does not test whether the second mutation produces an additional effect beyond the first. A separate epistasis-corrected analysis, in which multiple mutants were compared to the relevant single-mutant control, is presented in Supplementary Text Two.

### Supplementary Text 2

The standard training analysis in Dataset S9 reports an overall accuracy of 67 of 78 (86%), where each perturbation uses the baseline matched to its source publication. Among those 78 comparisons, 23 are pure multi-mutant perturbations (doubles or triples without exogenous treatment) and are compared with wild type following the convention used in the source literature. A true epistasis test instead asks whether stacking an additional mutation changes the phenotype beyond what the lower-order mutant background already produces, and so requires a different baseline. To carry out this test we re-derived the biological bin against the same lower-order mutant control by comparing the within-paper significance ranks reported in Dataset S2 (Bins sheet) for the multi-mutant and its first listed sub-mutant. A higher rank for the multi-mutant relative to its control gives a bin of 1, a lower rank gives a bin of −1, and the same rank gives a bin of 0. Per-paper bins were then aggregated across the supporting publications using the same strict two-thirds majority rule used in Dataset S4. The full per-paper derivation is provided as Dataset S17.

Of the 23 multi-mutant perturbations, 22 have a biological epistasis bin with consensus across the supporting papers, and one (D14_n, MAX4_n) does not and is listed alongside but excluded from the accuracy summary. On this 22-row analysis subset, PSoup matches the within-paper biological bin in 13 of 22 cases (59.1%) when scored under epistasis controls, compared with 19 of 22 (86.4%) under the standard WT controls (Table S8). This 27.3 percentage-point change isolates the model's ability to predict the additional effect of each successive mutation rather than the combined effect of all mutations against WT. Of the 22 comparisons, 10 were correct under both control schemes, 9 were correct only with WT controls, 3 were correct only with epistasis controls, and none were incorrect under both. The overall training accuracy of 67 of 78 (86%) is unchanged because this re-analysis only refines the interpretation of these multi-mutant rows.

Epistasis controls were chosen as the highest-order subtreatment that has a *Shoot Branching* value in Dataset S9 and is also reported in at least one source paper for the multi-mutant. For a double mutant A, B the control is typically the first listed single mutant A, and for a triple it is the highest-order subtreatment available in the same paper. The re-analysis is particularly informative for the strigolactone biosynthesis pathway. Under WT controls a double such as max3max4 appears correct because both the single and the double show increased branching relative to WT, but under epistasis controls the test asks whether the double shows more branching than the single, which the literature reports but the model often fails to predict. Table S8 lists the detailed outcome for all 23 perturbations, with rows that lack consensus shown for completeness and not scored.

Table S8. Detailed outcome for all 23 multi-mutant training comparisons under both wild-type and epistasis-corrected controls. Each multi-mutant is represented twice: compared with WT and compared with mutant for epistasis. The epistasis comparison uses the highest-order or lower-order-mutant control available in the source paper. For each multi-mutant, the baseline is the highest-order genotype reported in the source study that carries a subset of its perturbations. For a triple this is a double where one is available, otherwise a single. Where a double has both singles reported, the baseline is the single against which the source study draws its epistatic inference. Columns, left to right: Perturbation (the multi-mutant); Comparison with WT (the wild-type baseline); perturbation model output and WT model output (the simulated Shoot Branching values for the multi-mutant and wild type, with WT fixed at 1 by construction); Bins with WT, first the predicted (the bin from Pert. ÷ WT at the 5% threshold); and the Biological (the biological bin versus wild type, Dataset S4). The next section is the comparisons with mutant controls including the mutant baseline (the chosen epistasis baseline) and the model output for that mutant (its simulated Shoot Branching value); then the predicted bin value for the multi-perturbation compared with selected mutant baseline (epistasis; the bin from Pert. ÷ Mutant Baseline at the 5% threshold); and the Biological bin for this comparison (the within-paper rank-derived biological bin, Dataset S17). The final column in the match summery for comparison to WT and comparison for the epistasis using the baseline mutant (WT/Epi; the prediction is marked correct (✓) or incorrect (✗) under each scheme; A hyphen denotes a row that could not be scored, as no consensus biological expectation was available for that comparison.). Bins are coded as either an increase (+1), unchanged (0) or decrease (−1). Rows are sorted WT-only correct, then epistasis-only correct, then both correct, then both wrong, then the single no-consensus row (listed for completeness and not scored). Source data: Datasets S9, S4 and S17. Mutants are represented as perturbations to nodes written as GENE_n.

| | **Comparison with WT** | **Model Output** | | **Bins with WT** | | **Comparison with mutant** | **Model Output** | **Bins (Epistasis)** | | **Match** |
|---|---|---|---|---|---|---|---|---|---|---|
| **Perturbation** | **WT baseline** | **Pertub.** | **WT** | **Pred.** | **Biol.** | **Mutant baseline** | **Mutant** | **Pred.** | **Biol.** | **WT/Epi.** |
| BRC1_2_n, MAX_1_n | WT | 2.37 | 1.00 | +1 | +1 | BRC1_2_n | 2.00 | +1 | −1 | ✓ / ✗ |
| BRC1_2_n, MAX_2_n | WT | 2.37 | 1.00 | +1 | +1 | BRC1_2_n | 2.00 | +1 | −1 | ✓ / ✗ |
| BRC1_2_n, MAX_4_n | WT | 2.37 | 1.00 | +1 | +1 | BRC1_2_n | 2.00 | +1 | −1 | ✓ / ✗ |
| BRC1_2_n, PIN_1_3_4_7_n | WT | 1.25 | 1.00 | +1 | +1 | BRC1_2_n | 2.00 | −1 | 0 | ✓ / ✗ |
| D14_n, MAX3_n | WT | 1.27 | 1.00 | +1 | +1 | D14_n | 1.27 | 0 | +1 | ✓ / ✗ |
| D27_n, MAX4_n | WT | 1.27 | 1.00 | +1 | +1 | D27_n | 1.27 | 0 | +1 | ✓ / ✗ |
| MAX2_n, MAX1_n | WT | 1.27 | 1.00 | +1 | +1 | MAX2_n | 1.27 | 0 | +1 | ✓ / ✗ |
| MAX3_n, MAX4_n | WT | 1.27 | 1.00 | +1 | +1 | MAX3_n | 1.27 | 0 | +1 | ✓ / ✗ |
| SPL9_15_n, FHY3_FAR1_n | WT | 1.15 | 1.00 | +1 | +1 | SPL9_15_n | 1.33 | −1 | 0 | ✓ / ✗ |

| | | | | | | | | | | |
|---|---|---|---|---|---|---|---|---|---|---|
| PIN_1_3_4_7_n, MAX4_n | WT | 0.81 | 1.00 | −1 | +1 | PIN_1_3_4_7_n | 0.62 | +1 | +1 | ✗ / ✓ |
| SMXL6_7_8_n, MAX2_n | WT | 0.64 | 1.00 | −1 | 0 | MAX2_n | 1.27 | −1 | −1 | ✗ / ✓ |
| SMXL6_7_8_n, SPL9_15 | WT | 0.89 | 1.00 | −1 | 0 | SPL9_15_n | 1.33 | −1 | −1 | ✗ / ✓ |
| BRC1_2_n, D14_n | WT | 2.37 | 1.00 | +1 | +1 | BRC1_2_n | 2.00 | +1 | +1 | ✓ / ✓ |
| BRC1_2_n, HB21_40_53_n | WT | 2.00 | 1.00 | +1 | +1 | BRC1_2_n | 2.00 | 0 | 0 | ✓ / ✓ |
| BRC1_2_n, MAX4_n | WT | 2.37 | 1.00 | +1 | +1 | BRC1_2_n | 2.00 | +1 | +1 | ✓ / ✓ |
| BRC1_2_n, SMXL6_7_8_n, MAX2_n | WT | 1.38 | 1.00 | +1 | +1 | SMXL6_7_8_n, MAX2_n | 0.64 | +1 | +1 | ✓ / ✓ |
| D14_n, MAX2_n | WT | 1.27 | 1.00 | +1 | +1 | D14_n | 1.27 | 0 | 0 | ✓ / ✓ |
| D14_n, MAX3_n, MAX4_n | WT | 1.27 | 1.00 | +1 | +1 | D14_n, MAX4_n | 1.27 | 0 | 0 | ✓ / ✓ |
| D14_n, PIN_1_3_4_7_n | WT | 1.17 | 1.00 | +1 | +1 | D14_n | 1.27 | −1 | −1 | ✓ / ✓ |
| MAX2_n, MAX4_n | WT | 1.27 | 1.00 | +1 | +1 | MAX2_n | 1.27 | 0 | 0 | ✓ / ✓ |
| MAX4_n, MAX1_n | WT | 1.27 | 1.00 | +1 | +1 | MAX4_n | 1.27 | 0 | 0 | ✓ / ✓ |
| SMXL6_7_8_n, MAX3_n | WT | 0.64 | 1.00 | −1 | −1 | MAX3_n | 1.27 | −1 | −1 | ✓ / ✓ |
| D14_n, MAX4_n | WT | 1.27 | 1.00 | +1 | +1 | D14_n | 1.27 | 0 | no cons. | ✓ / - |

Of the 22 multi-mutant comparisons with biological epistasis consensus, 10 were correct under both control schemes, 9 were correct only with WT controls and represent cases where the model gets the WT-relative direction right but fails to predict the additional effect of stacking the second mutation, 3 were correct only with epistasis controls, and none were incorrect under both. The net effect is a loss of 6 correct predictions when switching from WT to epistasis controls, accounting for the change from 86.4% (19 of 22) to 59.1% (13 of 22). The cases where epistasis is not captured all lie within the same linear pathway. When one component is knocked out, the model cannot represent the additive effect of the second mutation, which indicates that an alternative pathway should be present in the model.

### Supplementary Text 3

To explore whether expanding sucrose signalling could improve model predictions we added three nodes to the original 32-node network, namely trehalose-6-phosphate (Tre6P), hexokinase 1 (HXK1) and the S1-bZIP transcription factor bZIP11, and connected them with ten new directed edges drawn from recent experimental evidence in Fichtner *et al.* (2021), Barbier *et al.* (2021), Kreisz *et al.* (2024) and Salam *et al.* (2021). The resulting network is shown in Fig. S3 and the raw results for the simulations in training and testing are in Dataset S14 and S15.

The expanded network contains 35 nodes and 66 edges. The original linear strigolactone biosynthesis chain from D27 through MAX3, MAX4, MAX1, CLAMT and LBO to SL was retained unchanged, and four existing equations were modified to incorporate the new sucrose signalling nodes. BRC1/2 gained Tre6P as an additional inhibitor, CK gained HXK1 as an additional stimulator, Bud_release gained Tre6P and HXK1 as stimulators and bZIP11 as an inhibitor, and SUC gained bZIP11 as an inhibitor to represent sugar export via SWEET transporters.

Eleven additional training perturbations were extracted from the sucrose signalling literature to test the new nodes. Three come from Fichtner *et al.* (2021) and test Tre6P overexpression and knockout, five come from Barbier *et al.* (2021) and test HXK1 knockout together with its epistatic interactions with cytokinin, decapitation and the strigolactone pathway, two come from Kreisz *et al.* (2024) and test bZIP11 knockout and overexpression, and one comes from Salam *et al.* (2021) and tests combined sucrose and cytokinin application. All eleven used the controls specified in the original publications, namely wild type for single mutants and single-mutant controls for double mutants. The perturbations, their simulated values under the 35-node network and the biological outcomes reported in each paper are listed in Table S10.

On the 78 original training perturbations the 35-node sucrose-integrated model reached an accuracy of 64 of 78 (82.05%), as reported in Dataset S14. On the eleven additional sucrose perturbations the model was correct in every case, giving 11 of 11 (100.0%, Table S9). Testing accuracy on the 84 testing pairs was 62 of 84 (73.81%), as reported in Dataset S15. The reduction in accuracy on the original training set relative to the 32-node base model (86% in Dataset S9) reflects the changed perturbation equilibria introduced by the additional sucrose nodes, since the perturbed steady-state values for nodes connected to the new sucrose edges (BRC1/2, CK, Bud_release and SUC) shift when mutations are introduced and this moves some comparison ratios across the 5% threshold. The fact that the sucrose-only perturbations were predicted correctly indicates that the added topology captures the core sucrose signalling logic, while the drop on the original set indicates that further tuning of the sucrose-connected equations is needed before the expanded network improves overall accuracy. The perturbations where the sucrose-integrated model disagreed with the biology are listed in Table S9.

Table S9. Eleven sucrose-signalling perturbations used to validate the 35-node sucrose-integrated network. The 11 perturbations test the added Tre6P, HXK1 and bZIP11 nodes (three from Fichtner *et al*. (2021); five from Barbier *et al*. (2021); two from Kreisz *et al*. (2024); one from Salam *et al*. (2021)). Columns, left to right: Perturbation; Compare with (its baseline); Pert and Compare (the simulated Shoot Branching values for the perturbation and the baseline, produced by the 35-node model); Pred. bin and Bio. bin (the predicted and biological directions of change, coded increase (+1), unchanged (0) or decrease (−1)); Match (✓ when the bins agree and ✗ otherwise); and Supporting paper. All 11 match the literature (11 of 11, 100%). Source data: Dataset S14.

| **Phenotypic Comparison** | | **Model Output** | | **Bins** | | | |
|---|---|---|---|---|---|---|---|
| **Perturbation** | **Compare with** | **Pert** | **Compare** | **Pred. bin** | **Bio. bin** | **Match** | **Supporting paper** |
| Tre6P OE (otsA) | WT | 1.80 | 1.00 | +1 | +1 | ✓ | (Fichtner *et al*., 2021) |
| Tre6P KO | WT | 0.44 | 1.00 | −1 | −1 | ✓ | (Fichtner *et al*., 2021) |
| brc1 x Tre6P OE | brc1 | 2.60 | 1.50 | +1 | +1 | ✓ | (Fichtner *et al*., 2021) |
| HXK1 KO (gin2) | WT | 0.57 | 1.00 | −1 | −1 | ✓ | (Barbier *et al*., 2021) |
| gin2 + decap | decap | 0.00 | 2.66 | −1 | −1 | ✓ | (Barbier *et al*., 2021) |
| gin2 + CK | gin2 | 0.89 | 0.57 | +1 | +1 | ✓ | (Barbier *et al*., 2021) |
| gin2/max4 | gin2 | 0.64 | 0.57 | +1 | +1 | ✓ | (Barbier *et al*., 2021) |
| gin2/max2 | gin2 | 0.64 | 0.57 | +1 | +1 | ✓ | (Barbier *et al*., 2021) |
| bZIP11 KO | WT | 4.12 | 1.00 | +1 | +1 | ✓ | (Kreisz *et al*., 2024) |
| bZIP11 OE | WT | 0.54 | 1.00 | −1 | −1 | ✓ | (Kreisz *et al*., 2024) |
| WT + SUC + CK | WT | 3.26 | 1.00 | +1 | +1 | ✓ | (Salam *et al*., 2021) |

Table S10. Sucrose-integrated training discrepancies (5% no-change band). The perturbations where the 35-node sucrose-integrated PSoup disagreed with the biology, of which nine are newly broken by the sucrose integration (correct under the base 32-node model) and the remainder were already wrong under the base model. Columns, left to right: Perturbation; Control (its baseline); Control branching and Pred. branching (the simulated Shoot Branching values for the control and the perturbation, produced by the 35-node model); Pred. bin and Bio. bin (the predicted and biological directions of change, coded increase (+1), unchanged (0) or decrease (−1)); Species; and Supporting Publication(s). Source data: Dataset S14.

| **Phenotypic Comparison** | | **Model Output** | | **Bins** | | | |
|---|---|---|---|---|---|---|---|
| **Perturbation** | **Control** | **Control branching** | **Pred. branching** | **Pred. bin** | **Bio. bin** | **Species** | **Supporting Publication(s)** |
| BRC1_2_n, +Low_R_FR | BRC1_2_n | 1.50 | 1.47 | 0 | −1 | Arabidopsis | (González-Grandío *et al*., 2017) |
| BRC1_2_n, PIN_1_3_4_7_n | WT | 1.00 | 0.94 | −1 | +1 | Arabidopsis | (van Rongen *et al*., 2019) |
| BRC1_2_n, PIN_1_3_4_7_n, +SL | BRC1_2_n, PIN_1_3_4_7_n | 0.94 | 0.92 | 0 | −1 | Arabidopsis | (van Rongen *et al*., 2019) |
| BRC1_2_n,+ High_R_FR | BRC1_2_n | 1.50 | 1.57 | 0 | +1 | Arabidopsis | (Aguilar-Martínez *et al*., 2007) |
| D14_n, PIN_1_3_4_7_n | WT | 1.00 | 1.00 | 0 | +1 | Arabidopsis | (Bennett *et al*., 2016) |
| FHY3_FAR1_n, +Low_R_FR | FHY3_FAR1_n | 0.90 | 0.85 | −1 | 0 | Arabidopsis | (Xie *et al*., 2020) |
| HB21_40_53_n, +ABA | HB21_40_53_n | 1.20 | 1.00 | −1 | 0 | Arabidopsis | (González-Grandío *et al*., 2017) |
| PIN_1_3_4_7_n | WT | 1.00 | 0.62 | −1 | 0 | Arabidopsis | (Bennett *et al*., 2016) |
| PIN_1_3_4_7_n, +SL | PIN_1_3_4_7_n | 0.62 | 0.61 | 0 | −1 | Arabidopsis | (van Rongen *et al*., 2019) |
| PIN_1_3_4_7_n, MAX4_n | WT | 1.00 | 0.70 | −1 | +1 | Arabidopsis | (van Rongen *et al*., 2019) |
| SMXL6_7_8_n, MAX2_n | WT | 1.00 | 0.84 | −1 | 0 | Arabidopsis | (Seale *et al*., 2017) |
| SPL9_15_n, +SL | SPL9_15_n | 1.20 | 1.17 | 0 | −1 | Arabidopsis, Rice | (Bennett *et al*., 2016) |
| WT, +NPA | WT | 1.00 | 0.57 | −1 | +1 | Rice | (Lin *et al*., 2009) |
| WT, +SL | WT | 1.00 | 0.97 | 0 | −1 | Arabidopsis, Pea, Petunia, Rice, Switchgrass | (Bennett *et al*., 2016) |

### Supplementary Text 4

#### Randomised-network comparison

To test whether the accuracy of the curated network depends on how its nodes are wired together, the same perturbations were simulated in randomised networks and scored exactly as the shoot branching network was scored. By wiring we mean which node connects to which, and the sign and influence type of each connection. No randomisation added or removed a node. Every randomised network contained the same 32 nodes as the curated network and the same logical rules, including AND conjunctions and necessary stimulation, which are properties of the model class rather than of the wiring (Fortuna *et al.*, 2026). A randomised network therefore differs from the curated network in its edges alone, which keeps the comparison fair because each randomised network remains the same class of model and differs only in the wiring being tested (Iorio *et al.*, 2016). AND groups were held together as single regulatory statements throughout, so that randomisation could not split a conjunction and thereby change the logic as well as the wiring.

Four randomisation schemes were used, ordered here from the scheme that preserves most of the curated structure to the scheme that preserves least. i) Sign shuffling keeps every edge where it is and permutes the stimulation and inhibition labels across the edges that carry them, so it tests whether the accuracy depends on the signs once the connections are correctly placed; necessary stimulation is exempt, because a necessary input is a structural statement rather than a sign. ii) Signed-degree-preserving rewiring moves edges but keeps, for each node, its number of incoming and outgoing edges and its balance of activating and inhibiting connections (Iorio *et al.*, 2016), so a randomised network resembles the curated one as closely as possible while still being randomised in its wiring; failure of this scheme is the strongest evidence that the specific arrangement of connections carries the regulatory logic. iii) Degree-preserving rewiring keeps only the number of incoming and outgoing edges per node (Maslov & Sneppen, 2002), so failure here shows that the accuracy is not explained by the degree distribution. iv) Erdős–Rényi rewiring keeps only the nodes and the number of edges of each influence type and joins them at random, giving the accuracy expected when almost no structure is retained, and is reported as a lower anchor rather than as evidence on its own.

Each scheme was run 1000 times against both the training and the testing responses (Ortiz-Gutiérrez *et al.*, 2015). Every randomised network was scored on whether the direction of change between the perturbation and its baseline matched the observed direction, using the same 5% no-change band applied to the curated model. For each scheme we counted how many randomised networks scored at least as well as the curated network and converted that count into an empirical P value (North *et al.*, 2002; Phipson & Smyth, 2010), calculated as $(1 + b) / (1 + m)$, where b is the number of randomised networks whose accuracy equalled or exceeded that of the curated network and m is the total number of randomised networks generated for that scheme, here 1000. With 1000 replicates the smallest attainable value is 1 / 1001, so a report of $P < 0.001$ means that no randomised network matched or exceeded the accuracy of the curated network. Randomised networks that did not reach a steady state were counted as failures rather than removed, because excluding them would condition the comparison on behaving like a real signalling network, which is the property under test. Results are given in Fig. S4 and Table S11.

#### What a randomisation actually changes

Because the schemes differ in what they disturb, it is worth being precise about what counts as a change. The curated network contains 52 regulatory statements. Counted as individual arcs the diagram has 56, because three statements are AND conjunctions with more than one input (D14, MAX2 and strigolactone together on strigolactone perception; BRC1/2 and low R:FR on HB21/40/53; gibberellin and polar auxin transport on shoot branching); these are held together throughout as single statements, so that randomisation cannot split a conjunction and thereby change the logic as well as the wiring. Of the 52 statements, 22 are inhibitions, 22 are stimulations and 8 are necessary stimulations.

After randomisation each of the 52 statements is in exactly one of three states, and the three always sum to 52 (Table S13). A statement is unchanged when it still runs between the same two nodes and still carries the same sign. It is sign-changed when it runs between the same two nodes but its sign has been inverted, so the connection is in the right place but acts the wrong way. It is moved when it now runs between a different pair of nodes. The distinction matters for sign shuffling in particular: that scheme moves nothing at all, yet leaves only a mean of 30.1 of the 52 statements unchanged, because 21.9 have had their sign inverted. The 30.1 that survive do so partly because permuting the labels returns some of them to their own statement by chance, and partly because the 8 necessary stimulations are exempt from shuffling: a necessary input is a structural statement, in that losing it drives the target to zero, rather than a sign. For the three rewiring schemes almost everything is moved rather than sign-changed, leaving a mean of 8.2, 5.3 and 0.9 statements unchanged respectively.

#### Whether fitting the training data transfers to the testing data

The testing set measures different nodes from the training set, bud release and transcript abundance rather than whole-plant branching, and so provides an independent check on whether a network fits the training perturbations for the right reasons. Randomised networks do not score low on the testing set in absolute terms; several schemes score higher there than on training, because the testing set has a different class balance (Table S11). What distinguishes the curated network is that its performance transfers. Selecting the ten best-fitting replicates of each scheme on training accuracy alone, accuracy fell by 20.7 percentage points between training and testing under sign shuffling, 28.2 under signed-degree-preserving rewiring and 37.3 under degree-preserving rewiring, against 11 for the curated network. Erdős–Rényi rewiring fell by 27.7 points, but only because its best replicates reach 59.2% on training and so have less to lose; for that reason the raw drop is reported here but not relied on. Within each ensemble, training and testing accuracy were only weakly correlated (Pearson r = 0.07 to 0.21), so a randomised network that fitted the training data was little more likely than any other to predict the held-out perturbations.

Comparing networks matched on how well they fit the training data removes that confound. Among replicates scoring between 70% and 80% on training, mean testing accuracy was 52.7% for sign shuffling, 46.0% for signed-degree-preserving rewiring and 37.7% for degree-preserving rewiring, and no Erdős–Rényi replicate reached that band at all. The same ordering holds in the 65% to 75% band (50.0%, 43.4% and 39.7%). Testing accuracy at equal training fit therefore tracks how much of the curated wiring each scheme leaves unchanged, a mean of 30.1, 8.2, 5.3 and 0.9 of the 52 statements respectively (Table S13). Sign shuffling, the only scheme that leaves every connection where it is and permutes only the signs, both fits and generalises best among the randomisations, and schemes that move the connections themselves generalise progressively worse. Across all 4000 randomised networks none reached the accuracy of the curated network on both datasets, and only one came within 10 percentage points on both (Fig. S5c, d).

A second and independent point is how rarely a randomised network fits the training data well in the first place. Reaching 70% training accuracy, still 16 percentage points below the curated network, required 58 of 1000 replicates under sign shuffling, 24 of 1000 under signed-degree-preserving rewiring and 14 of 1000 under degree-preserving rewiring, and no Erdős–Rényi replicate managed it at all (Table S13). This frequency tracks the same ordering as everything else, falling as the scheme leaves less of the curated wiring unchanged. The two observations close off complementary explanations: a randomised network rarely fits the training perturbations, and on the uncommon occasions when it does, that fit does not transfer to the held-out perturbations.

#### How far each randomised network sits from the curated one

Because a randomisation that rearranges almost every edge is a weaker test than one that rearranges a few, we measured the distance of every replicate from the curated network. Each replicate was regenerated from its original random seed and compared with the curated network edge by edge, counting the edges left in place, the edges displaced and the signs changed. The curated network contains 52 edges once AND groups are counted as single statements. Sign shuffling leaves all 52 edges in place and changes a mean of 21.9 signs. The three rewiring schemes are far more disruptive: degree-preserving rewiring leaves a mean of 5.3 of the 52 edges in place, signed-degree-preserving rewiring 8.2, and Erdős–Rényi rewiring 0.9. Across all 1000 degree-preserving replicates the single best-preserved network still retained only 12 of the 52 edges. This is a consequence of the standard mixing budget of ten times the number of edges used for this class of null model, and it means that the comparison in Fig. S4 establishes only that heavily rewired networks fail to reproduce the accuracy of the curated network. Within these ensembles the distance from the curated network is only weakly related to accuracy (Spearman rho between -0.02 and 0.25 across schemes and datasets), as expected when every replicate has already lost most of its curated wiring.

##### Networks that differ from the curated one by only a few edges

The near-curated regime was therefore tested directly, by displacing edges in controlled numbers rather than randomising the whole network. Using the degree-preserving swap with a budget of $k = 1, 2, 3, 5, 10, 20$ and 50 swaps and 200 replicates per value of k, mean training accuracy fell from 86% for the curated network to 74.8% after a single swap, which displaces two edges and leaves the other 50 in place, and then to 63.9%, 59.0%, 52.4%, 41.9%, 38.1% and 38.2% as more edges moved (Fig. S5b). At $k = 50$ the accuracy matches the fully mixed degree-preserving ensemble in Table S11 (37.3% training, 40% testing), confirming that the ladder and the published ensembles are measuring the same quantity. The spread at each value of k is wide: a single swap produced networks scoring anywhere between 20.5% and 89.7% on training, so the number of edges moved predicts accuracy only loosely and which edges move matters more than how many.

To characterise this exhaustively rather than by sampling, every admissible single swap was enumerated and simulated. Of the 1326 possible pairs of regulatory statements, 1016 are admissible once swaps that would create a self-loop, duplicate an existing edge or share a target are rejected. Displacing every edge cost accuracy on average, with no edge costing zero, so no edge in the curated network is redundant with respect to the data (Table S12). The most costly edges to displace were the necessary stimulation of shoot branching by bud release (53.6 points of training accuracy), the stimulation of bud release by cytokinin (26.5 points), the necessary stimulation of strigolactone perception by D14, MAX2 and strigolactone together (25.9 points) and the stimulation of cytokinin by SMXL6/7/8 (25.8 points). The least costly were the inhibition of MAX2 by sucrose (1.6 points) and the inhibition of strigolactone by CXE (4.1 points), the latter involving a node that no comparison in either dataset perturbs or measures.

A minority of these near-curated networks scored higher than the curated network: 110 of 1016 (10.8%) on training, 136 (13.4%) on testing and 27 (2.7%) on both. The gains were small, at most three additional training comparisons (70 of 78 against 67 of 78) and four additional testing comparisons (67 of 84 against 63 of 84), and they largely failed to generalise: of the 110 displacements that improved training accuracy, only 27 also improved testing accuracy while 46 made it worse. The comparisons these networks corrected were concentrated in two categories that the manuscript already attributes to the benchmark rather than to the topology. On the training set the 27 swaps corrected 40 comparisons in total, and 26 of the 40 were a single comparison, strigolactone applied to the brc1/2 pin1/3/4/7 double mutant and scored against that double mutant; a further 11 were multi-mutant genotypes scored against the wild type rather than against the appropriate lower-order mutant, the reference-genotype effect quantified in Table S8. On the testing set, 44 of the 56 corrections were measurements of MAX4 or SMXL6/7/8 transcript abundance, the readouts affected by the absence of feedback from strigolactone signalling onto strigolactone biosynthesis. The higher-scoring networks were also biologically implausible: the two that score highest on the training set (89.7%) require, in one case, phytochrome B to inhibit SMXL6/7/8 transcription while SMXL6/7/8 inhibits BRC1/2, and in the other, strigolactone perception to inhibit phytochrome B while low R:FR inhibits MAX4. Directional accuracy is therefore a necessary but not a sufficient criterion for a network of this kind. It constrains the wiring without uniquely determining it, and the curated topology is justified by the published evidence behind each edge rather than by being the highest-scoring arrangement of edges.

Table S11. Null-network comparison. For each randomisation family and each dataset: the accuracy of the curated topology, the mean and standard deviation of the null distribution over 1000 randomised networks, the range, the number of randomised networks scoring at least as well as the curated topology, the empirical P value computed as (1 + that number) / (1 + 1000) following North *et al*. (2002), and the proportion of runs reaching a steady state.

| **Randomisation** | **Set** | **Curated** | **Null mean ± SD** | **Null range** | **N ≥ curated** | **P** | **Steady state** |
|---|---|---|---|---|---|---|---|
| Erdős–Rényi rewiring | Training | 86% | 25.7 ± 11.7% | 2.6–62.8% | 0 / 1000 | < 0.001 | 87.2% |
| Erdős–Rényi rewiring | Testing | 75% | 28.2 ± 6.7% | 2.4–52.4% | 0 / 1000 | < 0.001 | 87.6% |
| Sign shuffling | Training | 86% | 37.8 ± 18.4% | 10.3–83.3% | 0 / 1000 | < 0.001 | 88.5% |
| Sign shuffling | Testing | 75% | 48.0 ± 7.9% | 28.6–71.4% | 0 / 1000 | < 0.001 | 86.9% |
| Degree-preserving rewiring | Training | 86% | 37.3 ± 13.4% | 10.3–78.2% | 0 / 1000 | < 0.001 | 92.4% |
| Degree-preserving rewiring | Testing | 75% | 40.0 ± 7.0% | 14.3–61.9% | 0 / 1000 | < 0.001 | 90.9% |
| Signed-degree-preserving rewiring | Training | 86% | 34.1 ± 14.3% | 9.0–79.5% | 0 / 1000 | < 0.001 | 88.2% |
| Signed-degree-preserving rewiring | Testing | 75% | 40.8 ± 7.7% | 11.9–65.5% | 0 / 1000 | < 0.001 | 87.6% |

Table S12. Per-edge accuracy cost from the exhaustive single-swap scan. Every one of the 1016 admissible single swaps of the curated network was simulated and scored; each swap displaces two edges, and the values below aggregate over all swaps that displace a given edge. Columns, left to right: Edge (source to target, with AND-grouped sources joined by +); Influence; Swaps (the number of single swaps that displace this edge); Training cost and Testing cost (the mean accuracy lost relative to the curated network's 86% and 75%, in percentage points); Worst training and Worst testing (the lowest accuracy observed across those swaps); and Region (whether either endpoint lies in the strigolactone or BRC1 core). Rows are sorted by training cost. No edge has a cost of zero, so no edge in the curated network is redundant with respect to the data. Source data: out/single_swap_scan.csv.

| Edge | Influence | Swaps | Training cost (pts) | Testing cost (pts) | Worst training (%) | Worst testing (%) | Region |
|---|---|---|---|---|---|---|---|
| Bud_release->Sustained_growth | necessary stimulation | 42 | 53.6 | 3.6 | 11.5 | 56.0 | periphery |
| CK->Bud_release | stimulation | 33 | 26.5 | 7.2 | 24.4 | 50.0 | periphery |
| SMXL6_7_8->CK | stimulation | 33 | 25.8 | 3.8 | 25.6 | 57.1 | SL/BRC1 core |
| D14+MAX2+SL->Perception_SL | necessary stimulation | 43 | 25.9 | 11.1 | 11.5 | 48.8 | SL/BRC1 core |
| Perception_SL->SMXL6_7_8 | inhibition | 35 | 22.8 | 10.8 | 34.6 | 58.3 | SL/BRC1 core |
| MAX1->CLAMT | necessary stimulation | 48 | 21.8 | 10.1 | 28.2 | 22.6 | SL/BRC1 core |
| CLAMT->LBO | necessary stimulation | 49 | 21.7 | 9.9 | 26.9 | 22.6 | SL/BRC1 core |
| LBO->SL | necessary stimulation | 48 | 21.3 | 8.0 | 24.4 | 22.6 | SL/BRC1 core |
| MAX4->MAX1 | necessary stimulation | 43 | 19.4 | 7.9 | 32.1 | 22.6 | SL/BRC1 core |
| NCED3->ABA | stimulation | 49 | 17.9 | 1.2 | 20.5 | 63.1 | periphery |
| BRC1_2+Low_R_FR->HB21_40_53 | stimulation | 45 | 16.7 | 1.5 | 20.5 | 63.1 | SL/BRC1 core |
| HB21_40_53->NCED3 | stimulation | 49 | 17.0 | 2.6 | 20.5 | 61.9 | periphery |
| BRC1_2->Bud_release | inhibition | 38 | 15.5 | 2.7 | 28.2 | 56.0 | SL/BRC1 core |
| MAX3->MAX4 | necessary stimulation | 38 | 13.7 | 4.8 | 46.2 | 44.0 | SL/BRC1 core |
| Decap_signal->Aux_Shoot | inhibition | 43 | 13.4 | 7.3 | 19.2 | 22.6 | periphery |
| Decap_signal->SUC | stimulation | 41 | 13.1 | 6.9 | 35.9 | 48.8 | periphery |
| GA+PAT_Bud->Sustained_growth | stimulation | 40 | 12.4 | 1.2 | 38.5 | 64.3 | periphery |
| SUC->Sustained_growth | stimulation | 34 | 12.8 | 1.8 | 28.2 | 64.3 | periphery |
| ABA->Bud_release | inhibition | 40 | 13.4 | 1.7 | 23.1 | 57.1 | periphery |

| | | | | | | | |
|---|---|---|---|---|---|---|---|
| SMXL6_7_8_Transcription->SMXL6_7_8 | stimulation | 42 | 12.0 | 1.1 | 15.4 | 65.5 | SL/BRC1 core |
| PhyB->MAX4 | inhibition | 32 | 11.6 | 4.2 | 17.9 | 42.9 | SL/BRC1 core |
| FHY3_FAR1->SMXL6_7_8 | stimulation | 40 | 11.3 | 4.1 | 33.3 | 60.7 | SL/BRC1 core |
| SPL9_15->BRC1_2 | stimulation | 35 | 10.6 | 5.5 | 23.1 | 57.1 | SL/BRC1 core |
| SMXL6_7_8->SMXL6_7_8_Transcription | inhibition | 43 | 10.6 | 5.6 | 12.8 | 58.3 | SL/BRC1 core |
| Aux_Shoot->CK | inhibition | 29 | 10.3 | 2.6 | 29.5 | 59.5 | periphery |
| SUC->Bud_release | stimulation | 33 | 10.0 | 2.4 | 29.5 | 57.1 | periphery |
| SMXL6_7_8->SPL9_15 | inhibition | 42 | 9.6 | 4.6 | 19.2 | 50.0 | SL/BRC1 core |
| CK->BRC1_2 | inhibition | 31 | 9.3 | 7.0 | 17.9 | 58.3 | SL/BRC1 core |
| PhyB->BRC1_2 | inhibition | 32 | 9.3 | 4.9 | 17.9 | 56.0 | SL/BRC1 core |
| SUC->CK | stimulation | 28 | 8.2 | 1.7 | 44.9 | 56.0 | periphery |
| PhyB->MAX2 | inhibition | 35 | 7.8 | 2.2 | 17.9 | 59.5 | SL/BRC1 core |
| Aux_Shoot->PAT_Bud | inhibition | 32 | 8.2 | 1.9 | 32.1 | 63.1 | periphery |
| Aux_Bud->GA | stimulation | 46 | 7.9 | 1.4 | 30.8 | 63.1 | periphery |
| D27->MAX3 | necessary stimulation | 42 | 7.4 | 3.5 | 20.5 | 40.5 | SL/BRC1 core |
| High_R_FR->PIFs | inhibition | 46 | 7.1 | 3.6 | 23.1 | 50.0 | periphery |
| PIFs->BRC1_2 | stimulation | 36 | 7.0 | 3.0 | 23.1 | 60.7 | SL/BRC1 core |
| Perception_SL->MAX4 | inhibition | 35 | 7.0 | 3.2 | 46.2 | 40.5 | SL/BRC1 core |
| Low_R_FR->FHY3_FAR1 | inhibition | 45 | 6.6 | 2.2 | 25.6 | 63.1 | periphery |
| CK->PAT_Bud | stimulation | 30 | 7.6 | 3.2 | 50.0 | 59.5 | periphery |
| Aux_Bud->PAT_Bud | stimulation | 39 | 7.0 | 1.4 | 29.5 | 63.1 | periphery |
| Decap_signal->PAT_Bud | stimulation | 39 | 7.0 | 1.7 | 29.5 | 63.1 | periphery |
| Aux_Shoot->MAX4 | stimulation | 32 | 5.8 | 3.9 | 21.8 | 44.0 | SL/BRC1 core |
| Aux_Shoot->MAX3 | stimulation | 35 | 5.7 | 3.5 | 21.8 | 44.0 | SL/BRC1 core |

| | | | | | | | |
|---|---|---|---|---|---|---|---|
| High_R_FR->FHY3_FAR1 | stimulation | 45 | 6.1 | 1.4 | 28.2 | 64.3 | periphery |
| FHY3_FAR1->SPL9_15 | inhibition | 42 | 5.7 | 2.3 | 20.5 | 56.0 | periphery |
| Perception_SL->MAX1 | inhibition | 38 | 5.6 | 2.8 | 44.9 | 40.5 | SL/BRC1 core |
| Perception_SL->MAX3 | inhibition | 37 | 5.6 | 1.3 | 46.2 | 61.9 | SL/BRC1 core |
| SUC->BRC1_2 | inhibition | 31 | 5.0 | 2.6 | 51.3 | 59.5 | SL/BRC1 core |
| Low_R_FR->PhyB | inhibition | 44 | 4.7 | 3.9 | 25.6 | 60.7 | periphery |
| High_R_FR->PhyB | stimulation | 44 | 4.5 | 2.0 | 28.2 | 63.1 | periphery |
| CXE->SL | inhibition | 49 | 4.1 | 0.6 | 23.1 | 63.1 | SL/BRC1 core |
| SUC->MAX2 | inhibition | 32 | 1.6 | 2.8 | 51.3 | 59.5 | SL/BRC1 core |

Table S13. What each randomisation scheme changes, and how often it fits the training data. The curated network contains 52 regulatory statements, with AND conjunctions counted once. After randomisation each statement is in exactly one of three states, and the three columns sum to 52. Columns, left to right: Scheme; Unchanged (the statement runs between the same two nodes and carries the same sign); Sign changed (same two nodes, inverted sign, so the connection is in the right place but acts the wrong way); Moved (the statement now runs between a different pair of nodes); Mean training and Mean testing accuracy across the 1000 replicates; Replicates reaching 70% training accuracy, out of 1000, which is still 16 percentage points below the curated network; and Mean testing accuracy of those replicates that fit the training data between 70% and 80%, which compares schemes at equal fit. Sign shuffling moves nothing yet leaves only 30.1 statements unchanged, because it inverts 21.9 signs; the 8 necessary stimulations are exempt from shuffling and are always unchanged. Reading down the table, the schemes that leave less of the curated wiring unchanged both fit the training data less often and generalise less well when they do. Curated network for comparison: 52 unchanged, 86% training, 75% testing. Source data: out/null_distances.csv and out/null_trials_*.csv.

| Scheme | Unchanged | Sign changed | Moved | Mean training (%) | Mean testing (%) | Reach 70% training (of 1000) | Mean testing at 70–80% training (%) |
|---|---|---|---|---|---|---|---|
| Sign shuffle | 30.1 | 21.9 | 0.0 | 37.8 | 48.0 | 58 | 52.7 |
| Signed-degree-preserving rewiring | 8.2 | 0.8 | 42.9 | 34.1 | 40.8 | 24 | 46.0 |
| Degree-preserving rewiring | 5.3 | 2.2 | 44.5 | 37.3 | 40.0 | 14 | 37.7 |
| Erdős–Rényi rewiring | 0.9 | 1.5 | 49.6 | 25.7 | 28.2 | 0 | none reach it |

#### Supplementary Datasets

The following datasets are provided as separate files in https://github.com/CMits/Shoot_Branching_PSoup.git.

- Dataset S1: Literature review papers used for creation of the knowledge graph.
- Dataset S2: Literature review papers used for extracting perturbations for training.
- Dataset S3: Literature review papers used for extracting perturbations for testing.
- Dataset S4: Training data — biological consensus for perturbations.
- Dataset S5: Testing data — biological consensus for perturbations.
- Dataset S6: Exogenous supply values for PSoup during training of the final model.
- Dataset S7: Genotype values for running PSoup during training of the final model.
- Dataset S8: Results of PSoup for the final trained model (training set).
- Dataset S9: Comparison of simulated vs observed bins for each training perturbation.
- Dataset S10: Exogenous supply values for PSoup during testing of the final model.
- Dataset S11: Genotype values for running PSoup during testing of the final model.
- Dataset S12: Results of PSoup for the final trained model (testing set).
- Dataset S13: Comparison of simulated vs observed bins for each testing perturbation.
- Dataset S14: Sucrose-integrated training results.
- Dataset S15: Sucrose-integrated testing results.
- Dataset S16: Epistasis analysis (training).
- Dataset S17: Biological epistasis bins (per-paper rank comparison).
- Dataset S18: Original network in SBGN form.
